# Dimerization GAS2 mediates microtubule and F-actin crosslinking

**DOI:** 10.1101/2024.08.19.608523

**Authors:** Jiancheng An, Tsuyoshi Imasaki, Akihiro Narita, Shinsuke Niwa, Ryohei Sasaki, Tsukasa Makino, Ryo Nitta, Masahide Kikkawa

## Abstract

GAS2 was originally identified as a growth arrest-specific protein, and recent studies have revealed its involvement in multiple cellular processes. Its dual interaction with actin filaments and microtubules highlights its essential role in cytoskeletal organization, such as cell division, apoptosis, and possibly tumorigenesis. However, the structural bases by which GAS2 regulates cytoskeletal dynamics remain unclear. In this study, we present cryo-EM structures of the GAS2- CH3 domain in complex with F-actin at 2.8 Å resolution, representing the first type 3 CH domain structure bound to F-actin, confirming its actin-binding activity. We also provide the first near-atomic resolution cryo-EM structure of the GAS2-GAR domain bound to microtubules and identified conserved microtubule-binding residues. Our biochemical experiments show that GAS2 promotes microtubule nucleation and polymerization and its C-terminal region is essential for dimerization, bundling of both F-actin and microtubules, and microtubule nucleation. Based on these results, we propose how GAS2 controls cytoskeletal organization.

## Introduction

The interplay between actin and microtubules (MTs) is crucial for maintaining cytoskeletal structures ^1–5^. Spectraplakin primarily mediates the interactions between actin filaments (F-actin) and microtubules (MTs), playing key roles in regulating cell polarity, cell-cell junctions, cell division, cell migration, and maintenance of neuron shape ^3,6,7^. However, the mechanisms and physical principles underlying spectraplakin-mediated actin–microtubule crosslinking are not well understood ^8^. No structural model currently explains these complex interactions.

Spectraplakins are a family of multi-domain proteins and include Dystonin (BPAG1), MACF1 (ACF7), and growth-arrest-specific 2 (GAS2) family, which comprises GAS2 and GAS2-like proteins 1-3 ^9–11^. Most spectraplakins contain two N-terminal calponin homology (C.H.) domains for F-actin binding, plakin and spectrin repeats, and a C-terminal GAR domain for microtubule binding (Supplementary Fig. 1a) ^9,12^. The GAS2 family crosslinks F-actin and MTs, which is essential for the regulation of axon morphology, cell apoptosis, and cell division ^7,13–15^. Notably, the GAS2 protein is the most diminutive spectraplakin, comprising a single type 3 CH domain and a GAR domain connected by a hinge linker (H-Linker, Fig. 1a, first panel).

**Fig. 1.**
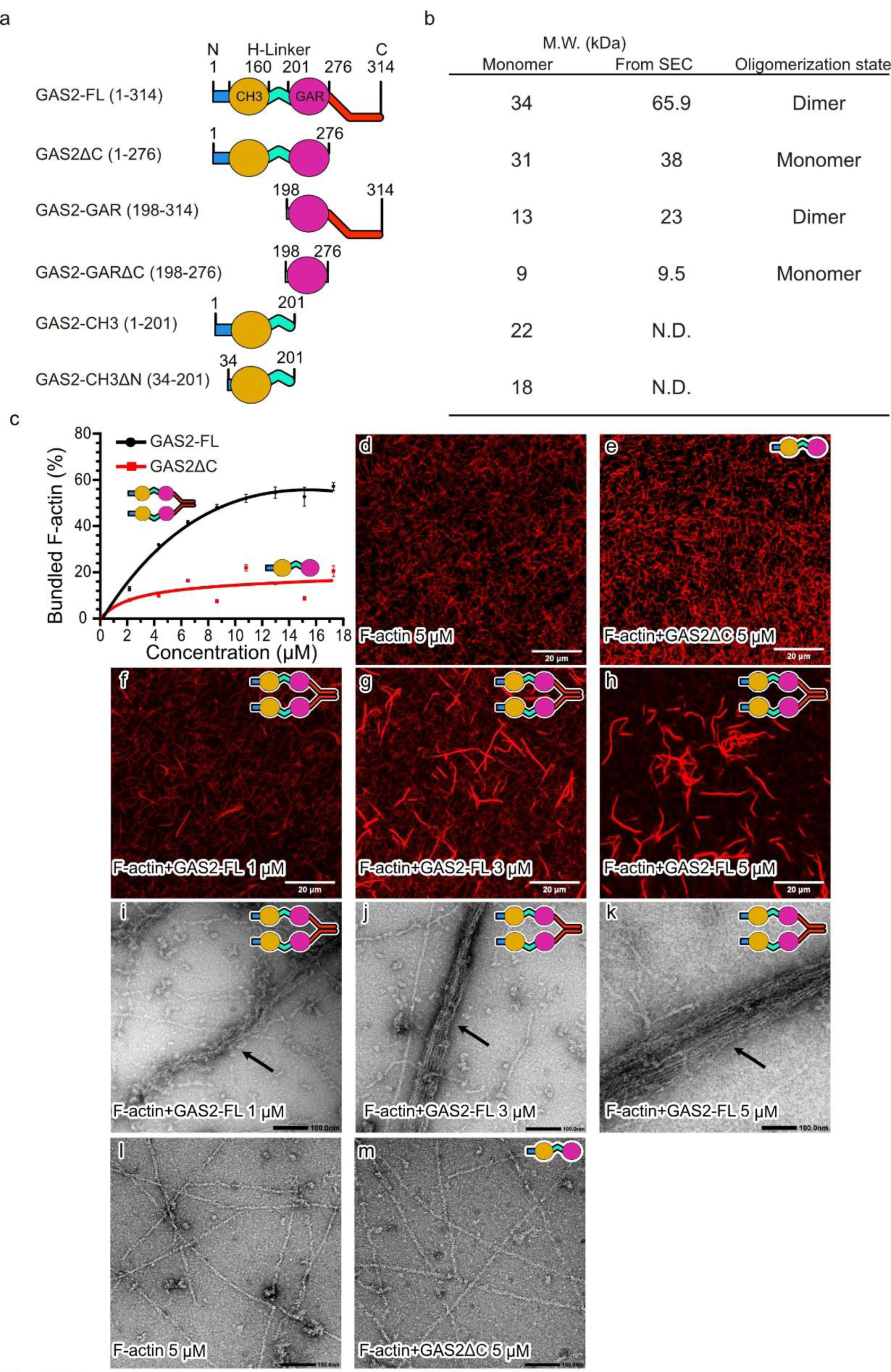
The C-terminal of GAS2 is necessary for dimerization and is essential for F-actin bundling. a. GAS2 constructs used in biochemical assays. The first panel shows that the full- length GAS2 (GAS2-FL) consists of the CH3 and GAR domains, which are connected by a hinge linker (H-Linker). Mouse GAS2-FL is a 314 amino acids protein, and the theoretical monomer M.W. (molecular weight) is about 34 kDa. The second panel shows the GAS2ΔC (residues 1-276), the theoretical monomer M.W. is approximately 31 kDa. The third panel shows the GAS2-GAR domain (residues 198-314), the theoretical monomer M.W. is about 13 kDa. The fourth panel shows the GAS2-GARΔC (residues 198-276), the theoretical monomer M.W. is about 9 kDa. The fifth panel shows the GAS2-CH3 domain (residues 1- 201). The sixth panel shows the GAS2-CH3ΔN (residues 34-201). **b** GAS2 constructs theoretical M.W., computational M.W. of the SEC (Size Exclusion Chromatography) results, and oligomerization state. **c** Dose-dependent bundling curve of F-actin by GAS2-FL or GAS2ΔC for low-speed co-sedimentation assay. F-actin (5 μM as monomer) was incubated with increasing concentrations of GAS2-FL or GAS2ΔC (2–16 μM as monomer) and actin- bundling assays were performed. The amount of bundled F-actin was plotted against the concentration of GAS2-FL or GAS2ΔC for SDS-PAGE results in (**supplementary** Fig. 1h**- i**). Mean±SD. from three independent experiments is shown. **d-h** Confocal fluorescence microscopy bundle assay in the presence or absence of GAS2-FL or GAS2ΔC. Actin filaments were labeled by Alexa Fluor 568 phalloidin and imaged under confocal microscopy. **i-m** Representative negatively stained electron micrographs of F-actin in the presence or absence of purified of GAS2-FL or GAS2ΔC. Black arrows show the F-actin bundle arrays.

The CH domain, a ∼110-residues protein module, is known for its roles in actin-binding, signaling, and microtubule binding. The CH domains are highly conserved, with four major helices forming a compact globular structure ^16^. Three major groups of CH domain have been recognized—CH1, CH2, and CH3—based on sequence alignments and variations in loop or short helix structures ^17,18^. CH1 domains typically have a longer first major helix; the CH2 family features a short helix following the second long major helix, a structure absent in CH1 and CH3; the CH3 domain uniquely contains a short four-residue 3^10^ helix after the third main helix ^19^. CH1 and CH2 domains appear as tandem pairs in MACF1, BPAG1, and Plectin, performing actin binding and bundling functions^18,20^. In contrast, single CH3 domains are found in proteins involved in muscle contraction regulation, such as calponin, and in signal-transduction proteins like Vav and IQGAPs. The interaction of the single CH3 domain with F-actin remains controversial ^16,21–23^. The differences in F-actin binding among CH1 and CH3 domains are not well understood. Additionally, a special group of single CH domains is present in EB proteins (EB-type CH domain), which specifically bind to the microtubule plus end^24^. The mechanisms by which this highly conserved CH domain recognizes the difference between the microtubules and F-actin remain elusive.

The GAR domain consists of approximately 70 amino acids and binds to the MT lattice ^25–28^. The GAR domain is highly conserved from *C. elegans* to humans. According to an X-ray crystallographic analysis of the GAR domain of ACF7, it has a unique zinc-binding α/β fold, wherein zinc plays a pivotal role in maintaining the structural integrity of the domain ^29^. The GAR domain is essential for MACF1’s microtubule binding and is critical for neuronal development and axon guidance. Mutations in the zinc-binding motif of MACF1’s GAR domain disrupt this interaction, resulting in lissencephaly and brainstem hypoplasia ^30^.

GAS2, a component of the microfilament system, is processed by caspase enzymes during apoptosis, which cleaves its carboxyl-terminal (C-terminal) domain. This cleavage leads to significant reorganization of the microfilament system, altering cell shape and playing a crucial role in the morphological changes associated with cell death ^31^. However, the function of the C-terminal is always elusive. Additionally, during the G0 to G1 transition of the cell cycle, GAS2 is rapidly phosphorylated in response to growth factors, resulting in its relocation to membrane ruffles at cell edges and inducing significant microfilament rearrangement ^13,32^. In vitro studies further suggest that Gas2 is involved in the differentiation of keratinocytes and muscle cells, and can inhibit cell division, resulting in multinucleated cells by microtubule bundling ability ^15^. In vivo studies demonstrate that GAS2 functions as a death substrate in apoptosis, influencing chondrocyte differentiation and limb myogenesis ^14^. Notably, GAS2 is also expressed in Pillar and Deiters’ cells, which are crucial for sound wave transduction to hair cells. These cells possess a rigid cytoskeleton composed of hundreds to thousands of microtubules organized into tightly bundled arrays cross-linked with actin filaments (Supplementary Fig. 1b) ^33^. GAS2 co-localizes with these microtubule bundles, and the variants (GAS2 c.723+1G>A; GAS2 c.616–2 A > G) introduce a stop codon within the GAR domain, resulting in disordered microtubule arrays and causing inherited hearing loss^33,34^. However, its molecular mechanism is poorly understood.

Although GAS2 is essential for cytoskeleton rearrangement during apoptosis, the mechanisms by which GAS2 mediates these changes are not well understood. The functional implications of specific GAS2 mutations causing hearing loss remain unclear due to a lack of structural data and limited functional characterization. To address these gaps, here, we obtained a cryo-EM structure of the F-actin complexed with the CH3 domain of GAS2 at 2.8 Å resolution and a cryo-EM structure of the GAS2-GAR domain decorated on microtubules at 3.2 Å resolution. Our co-sedimentation, negative staining, and TIRF assays demonstrate that GAS2 binds to F-actin, interacts with microtubules, and promotes microtubule nucleation and stabilization. Additionally, GAS2 bundles microtubules and F-actin at low concentrations, suggesting a role in maintaining cell stiffness. Our findings reveal that the C-terminal region of GAS2 is essential for its dimerization, tubulin nucleation, and bundling abilities, providing significant insights into the molecular mechanisms of GAS2-mediated F-actin to microtubule cross-linking. This may pave the way for new therapeutic strategies for diseases involving dysregulated microtubule and actin dynamics.

## Results

### The C-terminal of GAS2 is necessary for dimerization and is essential for F-actin bundling

Full-length GAS2 (GAS2-FL (1-314)) consists of the CH3 and GAR domains interconnected by a hinge linker (H-Linker, Fig. 1a). To evaluate the functional importance of each domain, we generated a series of deletion mutants and assessed them using biochemical assays, TIRF microscopy, and negative stain electron microscopy (EM) (Fig. 1). Analysis by size exclusion chromatography (SEC) showed that the peak location of GAS2-FL corresponds to 66 kDa, indicating that the protein forms a homodimer (Fig. 1a, b, Supplementary Fig. 1c-e). On the other hand, GAS2ΔC, lacking the C-terminal domain, appeared as a monomer protein (Fig. 1a, b, Supplementary Fig. 1c-e), suggesting that the C-terminal domain is important for dimerization. To further identify the minimum region required for dimerization, we generated GAS2-GAR domain constructs with and without the C-terminal region (GAS2-GARΔC) to further identify the minimum region for dimerization and performed SEC experiments (Fig. 1a). The results showed that GAS2-GAR is a dimer, while GAS2-GARΔC is a monomer (Fig. 1b, Supplementary Fig. 1c- e). These findings demonstrate the critical role of GAS2’s C-terminal domain in its dimerization.

We utilized a co-pelleting assay to investigate the interaction between full-length GAS2 and actin filaments in vitro. Both full-length GAS2 and its CH3 domain (Fig. 1a, fifth panel, amino acids 1- 201) were co-sedimented with actin filaments (Supplementary Fig. 1f, g), indicating that the CH3 domain is responsible for binding to F-actin. We performed a low-speed co-sedimentation assay to test whether GAS2 promotes F-actin bundling ^35^. F-actin alone does not sediment at low centrifugal forces (9,000 × g) (Supplementary Fig. 1h, leftmost lane). However, increasing concentrations of full-length GAS2 resulted in a concentration-dependent increase in the amount of actin in the pellet, indicating GAS2-induced F-actin bundling (Fig. 1c, Supplementary Fig. 1h).

In contrast, when GAS2ΔC was incubated with F-actin, the amount of actin in the pellet did not increase to the same extent as observed with full-length GAS2 (Fig. 1c, Supplementary Fig. 1i). At 16 μM concentration, full-length GAS2 bundled 60 ± 1.9% of F-actin, while GAS2ΔC only bundled 23 ± 2.6% under the same conditions (Fig. 1c). This suggests that the C-terminal domain of GAS2 is necessary for efficient F-actin bundling. Fluorescence microscopy further supported these findings; GAS2 addition led to longer, thicker filaments with enhanced fluorescence, indicating actin bundling (Fig. 1f-h). Conversely, the GAS2ΔC failed to induce significant changes (Fig. 1e), similar to the bare F-actin control (Fig 1d). Negative stain electron microscopy further confirmed these observations; in the absence of GAS2-FL, actin filaments remained disordered (Fig. 1l), whereas in its presence, actin filaments bundled into higher-order structures (Fig. 1i-k). GAS2ΔC failed to induce bundling (Fig. 1m).

In conclusion, our results prove that the CH3 domain of GAS2-FL can interact with actin filaments. The low-speed co-sedimentation assay demonstrated that GAS2-FL can bundle actin filament and that the C-terminal (residues 277-314) of GAS2 is essential for this activity.

### Cryo-EM structure of GAS2 CH domain bound to F-actin

To understand how GAS2 recognizes F-actin through its CH3 domain at the molecular level, we utilized cryo-EM single particle analysis. We used GAS2-CH3, the minimum construct that can bind to F-actin and obtained a high-quality density map with a resolution better than 3.0 Å, as judged by the FSC curve (Table 1, Fig. 2a, Supplementary Fig. 2a). To model the atomic structure, we used the AlphaFold2 prediction as a starting model (AF2 model No. AF-P11862-F1) ^36^. The AF2 model fits very well, except for a few flexible loops. After several rounds of manual modeling and refinement, we obtained the final atomic model (Fig. 2c, Supplementary Fig. 2b).

**Fig. 2.**
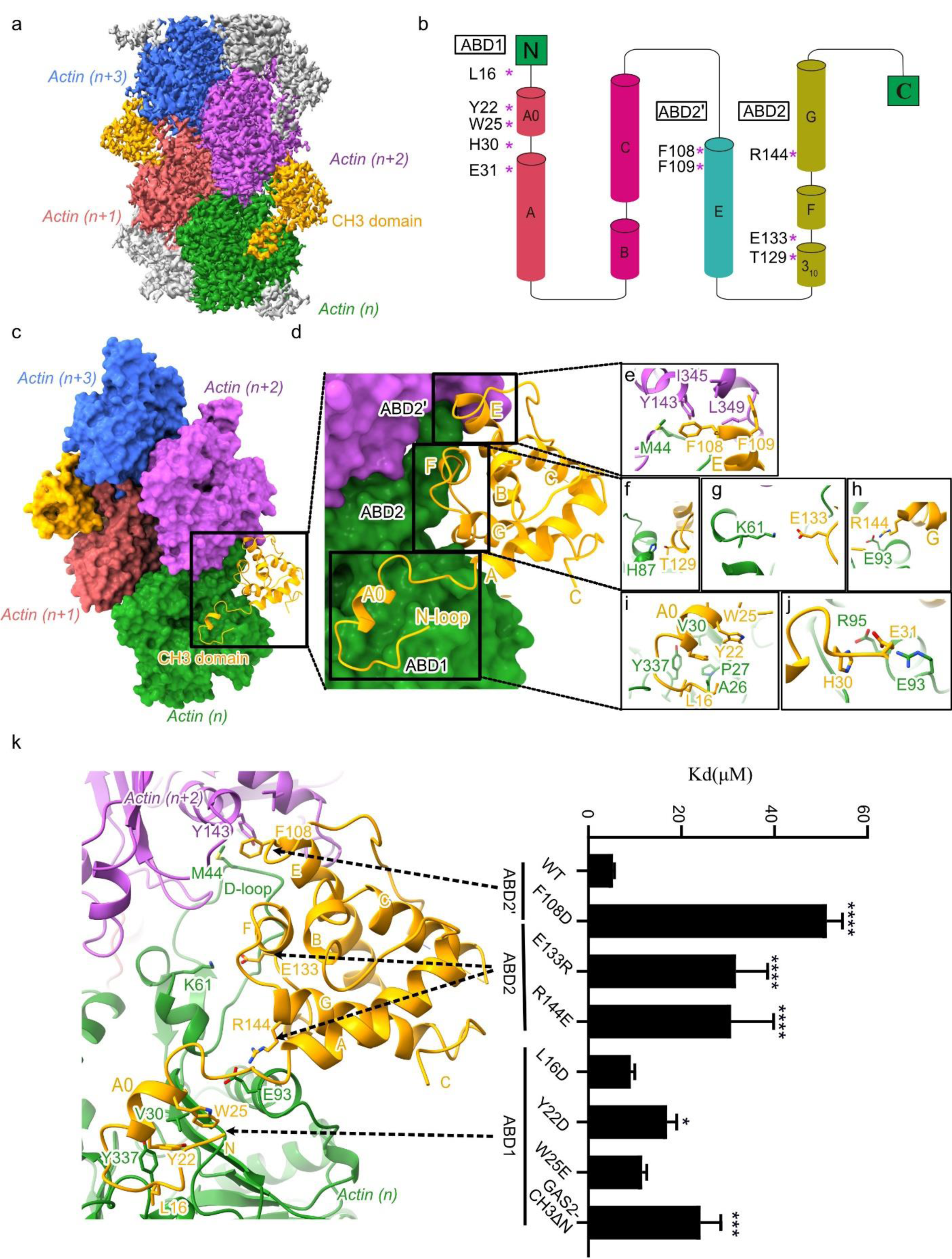
Cryo-EM map and model of GAS2-CH3 domain bound to F-actin. a. Reconstruction cryo-EM map of GAS2-CH3 domain decorated on F-actin at a 2.8 Å resolution. 4 Actin binding subunits (marked as n series from bottom to top) are presented in green, indian red, medium orchid, royal blue and GAS2-CH3 domain is presented in orange, respectively. **b** Topology structure of GAS2-CH3 domain and the main interaction amino acids in the topology structure. Four main helices labeled with A, C, E, G; short helices including A0, B, F and a short 3_10_ helix. **c** Atomic model of GAS2-CH3 domain decorated on F-actin with the same color scheme as in (**a)**, the GAS2-CH3 domain in orange interacts with two adjacent actin monomers thus following the actin helical pattern. **d** Closer view of GAS2-CH3-F-actin model, there are mainly three actin binds sites for GAS2-CH3 domain, respectively ABD1, ABD2’, and ABD2 are shown as in black box. **e-j** Residual level information of key amino acids interacting with actin monomers from ABD1: L16, Y22 and W25 in (**i)**, H30 and E31 in (**j)**; ABD2’: F108, F109 in (**e)**; ABD2: T129 in (**f)**, E133 in (**g),** and R144 in (**h)**. **k** The histogram of *K_d_* values for GAS2-CH3 WT and mutants of representative Coomassie-stained actin co-sedimentation assay. *K_d_* values are shown in (Supplementary Fig. 2c), and the fitting curve are shown in (Supplementary Fig. 2d). Raw SDS-PAGE gels are shown in source data. Bars = mean ± SD. from three independent experiments, * (p-value < 0.05), *** (p-value < 0.001), **** (p-value < 0.0001) significantly different from WT in a one-way ANOVA with Dunnett’s multiple comparisons test.

**Table 1.**

| Data collection and processing |  |  |  |
| --- | --- | --- | --- |
| Complex structure | GAS2-CH3-F-actin | GAS2-GAR-MT | GAS2-FL-MT |
| Magnification | 105,000 | 105,000 | 105,000 |
| Voltage (kV) | 300 | 300 | 300 |
| Electron exposure (e<sup>-</sup>/Å<sup>2</sup>) | 48.93 | 48.79 | 49.92 |
| Defocus range (μm) | -2.4 to -1.2 | -1.8 to -1 | -1.8 to -1 |
| Pixel size (Å) | 0.83 | 0.85 | 0.83 |
| Number of images taken | 6378 | 3322 | 261 |
| Symmetry imposed | Pseudo-Helical | Pseudo-Helical | Pseudo-Helical |
| Initial particles number | 632,290 | 70012 | 22,841 |
| Final particles number | 412,176 | 44,922 | 5,616 |
| Map resolution (Å) | 2.8 Å | 3.2 Å | 4.39 Å |
| FSC threshold | 0.143 | 0.143 | 0.143 |
| Map sharpening B factor (Å<sup>2</sup>) | -111 | -89 |  |
| Refinement |  |  |  |
| Initial model used | AlphaFold:<br>AF-P11862-F1 | PDB:1V5R |  |
| Model resolution (Å°)/FSC threshold | 2.83/0.5 | 3.28/0.5 |  |

| Model composition |  |  |
| --- | --- | --- |
| Non Hydrogen atoms | 13,802 | 7,434 |
| Protein residues | 1,756 | 937 |
| Ligands | MG:4/ADP:4 | GTP:2/ZN:1/MG:2 |
| R.m.s. deviations |  |  |
| Bond lengths (Å) | 0.029 | 0.005 |
| Bond angles (°) | 3.736 | 1.174 |
| Validation |  |  |
| MolProbity score | 1.98 | 2.41 |
| Clash score | 1.64 | 13.67 |
| Ramachandran plot |  |  |
| Favored (%) | 94.89 | 93.66 |
| Allowed (%) | 3.97 | 6.12 |
| Outliers (%) | 1.15 | 0.21 |

Our atomic model identified three main actin-binding sites in the GAS2-CH3 domain, namely ABD1 (amino acids 13-31), ABD2’ (amino acids 107-111), and ABD2 (amino acids 127-145) (Fig. 2d). At ABD1 (amino acids 13-31), the N-terminal loops or helixes of the GAS2-CH3 domain mainly interact with actin (n) by hydrophobic or electrostatic environment (Fig. 2b, d, i, j). At ABD2’, the n+2 position of actin Y143, I345, L349, and n position of actin M44 at the D-loop form a hydrophobic pocket that enfolds F108 and F109 of the GAS2-CH3 domain (Fig. 2b, d, e). ABD2 had electrostatic interactions with the n position of actin between actin H87 and GAS2 T129, actin K61 and GAS2 E133, actin E93 and GAS2 R144 (Fig. 2b, d, f, g, h).

To validate our structural model, we generated point mutation proteins and analyzed them using an F-actin co-pelleting assay. We introduced mutations (L16D, Y22D, W25E, and GAS2-CH3ΔN (Fig. 1a bottom panel)) targeting to disrupt ABD1 hydrophobic interactions, one mutation (F108D) targeting the hydrophobic pocket in ABD2, and two mutations (E133R and R144E) targeting electrostatic interaction in ABD2 (Fig. 2k, Supplementary Fig. 2c, d). Compared to the wild-type CH3 domain, Y22D, GAS2-CH3ΔN, F108D, E133R, and R144E showed significantly lower affinity for F-actin (Fig. 2k, Supplementary Fig. 2c, d). These findings support the conclusion that a single GAS2-CH3 domain is sufficient for interaction with F-actin. Furthermore, our study highlights the critical roles of hydrophobic pocket in ABD1, ABD2’, and ABD2 electrostatic interactions between the CH3 domain and F-actin.

### GAS2 bundles Microtubule by binding to MT through GAR domain

Utilizing co-pelleting assays, we confirmed that both full-length GAS2 and its GAR domain (198- 314) exhibit binding affinity to taxol-stabilized MTs (Supplementary Fig. 3a, b). Dark-field microscopy directly visualized MT bundling by different GAS2 constructs. We found that GAS2- FL and GAS2-GAR, particularly with an intact C-terminal region, induced dense clustering and bundling of MTs (video 1 for the bare MT; video 2 for GAS2-FL; video 4 for GAS2-GAR). In contrast, GAS2-ΔC and GAS2-GARΔC without the C-terminal failed to produce the effects (video 3 for GAS2ΔC; video 5 for GAS2-GARΔC). Electron microscopy further elucidated well-ordered MT bundles formed in the presence of GAS2-FL compared with bare MT (Fig. 3a, d-f), emphasizing the crucial role of the C-terminal region in MT bundling. Additionally, the GAS2- GAR domain demonstrated significant MT bundling ability (Fig. 3g-i). In contrast, GAS2ΔC and GAS2-GARΔC, which lack the C-terminal region, exhibited an inability to induce MT bundling (Fig. 3b, c). These findings underscore the essential role of the C-terminal region in GAS2- mediated MT bundling.

**Fig. 3.**
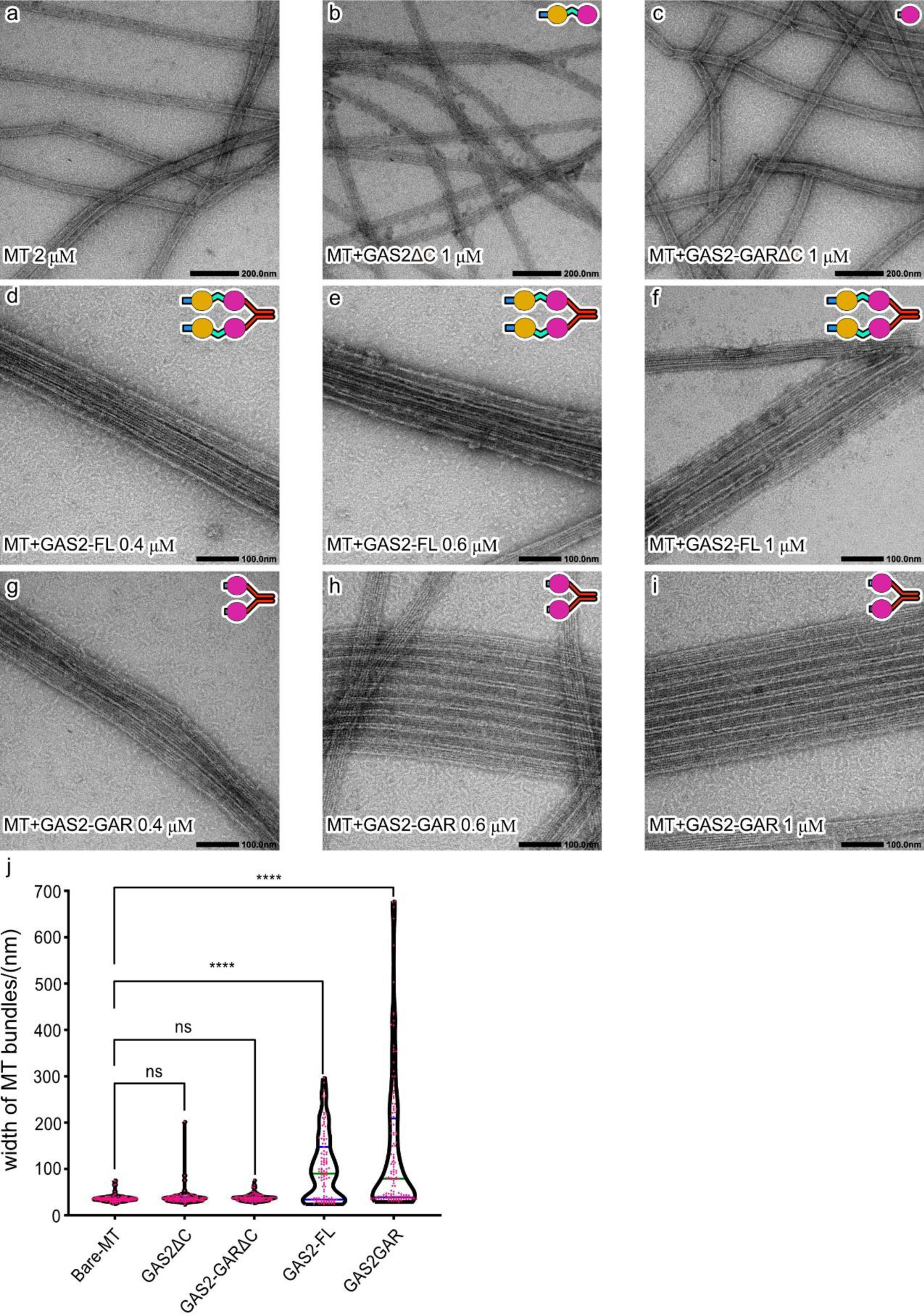
Negative staining electron microscopy reveals the GAS2 MTs bundling ability. a. Bare MT is disordered as individual filaments from negative staining micrographs. **b** GAS2ΔC did not bundle MT, and most of MTs are disordered individual filaments. **c** GAS2- GARΔC did not bundle MT and most of MTs are disordered individual. **d-f** GAS2-FL gathered MT to well-ordered bundles at different concentrations. **g-i** GAS2-GAR domain gathered MT to well-ordered bundles at different concentrations. **j** Width of MTs or MTs bundles were measured. In one micrograph, all MTs or MTs bundles width were measured. For control, bare-MT N=114 MTs or MTs bundles were checked; For GAS2ΔC N=111 MTs or MTs bundles were checked; For GAS2-GARΔC N=103 MTs or MTs bundles were checked; For GAS2-FL N=108 MTs or MTs bundles were checked; For GAS2-GAR N=137 MTs or MTs bundles were checked. One-way ANOVA with Dunnett’s multiple comparisons test has been used. ****, p-value < 0.0001, ns: no significant difference. All the detailed values are shown in Source data.

We measured the diameter of MTs and MT bundles in micrographs and found that the average diameter of bare MTs was about 30 nm. However, in the presence of GAS2-FL or GAS2GAR, the average width of MT bundles widened significantly to approximately 150 nm or 250 nm, respectively (Fig. 3j). Conversely, there was minimal change in MT diameter with GAS2ΔC or GAS2-GARΔC presence when these proteins lost the C-terminal (Fig 3j). In summary, our results furnish compelling evidence that the C-terminal region of GAS2 is pivotal for bundling MT.

### Cryo-EM structure of GAS2-GAR-microtubule complex

To further investigate the interaction between GAS2 and MT, we used the GAS2-GAR domain construct to elucidate the GAR structure bound to microtubules by single-particle cryo-EM analysis. After eliminating poorly decorated particles, 2D class averages exhibit GAS2-GAR domain distribution along microtubule surfaces with 8 nm intervals (Supplementary Fig. 3c, d).

We employed MiRP ver2 to analyze microtubule, which exhibits a pseudo-symmetric nature with a seam ^37^ . Our analysis achieved a 3.2 Å resolution of 14-pf microtubules complexed with GAS2- GAR domain (Supplementary Fig. 3c). Local resolution estimates varied due to conformational heterogeneity, ranging from 3.0 to 3.5 Å for tubulins and 3.5 to 4 Å for GAR domain molecules (Supplementary Fig. 3e).

Analysis of the density map reveals the insertion of the GAR domain into the tubulin intradimer groove (Fig. 4a, b). At the GAR-tubulin interface, the local resolution was approximately 3.5 Å (Supplementary Fig. 3e), enabling observation of amino acid sidechains. Notably, the resolutions of N- and C-termini and certain loops of the GAS2-GAR domain dip below 3.8 Å, possibly due to its flexibility (Supplementary Fig. 3e).

**Fig. 4.**
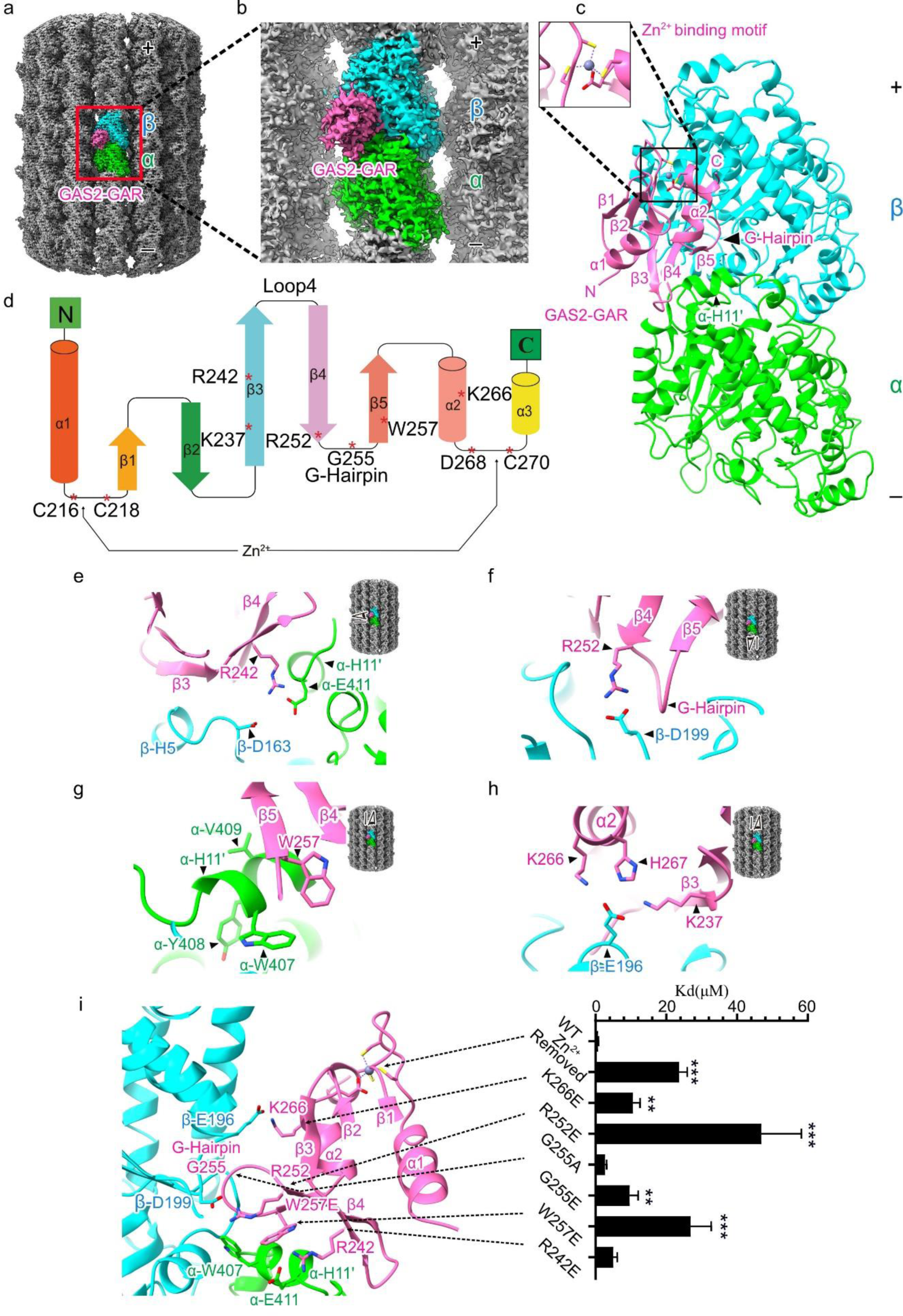
The 3.2-Å resolution cryo-EM structure of the GAS2-GAR domain bound to the microtubule. a. Cryo-EM map of 14-protofilament GMPCPP-stabilized microtubule decorated with GAS2-GAR domain. α-tubulin: lime, β-tubulin: cyan, and GAS2-GAR domain: hot-pink. **b** Enlarged view of the Cryo-EM map of the red box in (**a**). **c** Atomic model of GAS2-GAR-tubulin complex. The G-hairpin stench into intradimer of the tubulin. The zinc-binding motif of GAS2-GAR domain is shown in the enlarged view. **d** Topology structure of GAS2-GAR domain and the main interaction amino acids in secondary structure. All these interactions are located in β3-β4-β5-α2 regions and the loops between β3-β4-β5- α2. The C216, C218, D268 and C270 form the zinc binding motif. **e** The long β3 of the GAS2GAR domain spans from β-tubulin to α-tubulin. The R242 of the GAR domain is close to H11’ of α-tubulin’s E411, which indicates R242 may form a salt bridge with α-tubulin’s E411. And R242 is also close to β-tubulin D163 and potentially interacts with β-tubulin D163. **f** The G-hairpin penetrates into α-β tubulin intradimer. The R252 is close to β-tubulin D199 in the loop between H5 (helix 5) and S6 (sheet 6), which indicates that R252 may form a salt bridge with β-tubulin D199. **g** α-tubulin H11’ W407, Y408, V409, and G410 form a hydrophobic pocket. W257 in β5 of the GAR domain nestles into α-tubulin H11’ hydrophobic pocket and is close to α-tubulin W407. **h** K237 in β3, K266 and H267 in α-helix 2 of the GAS2-GAR domain is close to E196 in β-tubulin. **i** The histogram of *K_d_* values for GAS2- GAR WT and mutants of representative Coomassie-stained co-sedimentation assay. *K_d_* values are shown in (**Supplementary** Fig. 3h), and the fitting curves are shown in (**Supplementary** Fig. 3i). Raw SDS-PAGE gels are shown in source data. Bars = mean ± SD. from three independent experiments, ** (p-value < 0.01), *** (p-value < 0.001) significantly different from WT in a one-way ANOVA with Dunnett’s multiple comparisons test.

To elucidate MT-GAR atomic details of the GAS2-MT interaction, we modeled a GAS2-GAR- MT complex using the previously reported NMR structure (PDB: 1V5R, Supplementary Fig. 3f) and αβ-tubulin heterodimers (PDB: 3JAT) as starting models. After several rounds of modeling and refinement, we obtained the final model (Fig. 4c, Supplementary Fig. 3g). GAS2-GAR domain interaction with MT via β3-β4-β5-α2 regions and loops. Notably, β3 of the GAR domain forms hydrophilic interactions with β-tubulin H5 and α-tubulin H11’. Specifically, GAS2 R242 potentially engages in a salt bridge with α-tubulin E411 (Fig. 4e). Additionally, GAS2 R242 faces β-tubulin D163, possibly facilitating interaction (Fig. 4e). Similarly, GAS2 β4 R252 proximity to β-tubulin D199 implies a potential salt bridge consistent with the observed density (Fig. 4f).

Within the β4-β5 loop of the GAS2-GAR domain, there are three glycine residues (G254, G255, G256), which are highly conserved across GAR domain homologs. The β4-β5 loop form hairpin structure (G-hairpin) (Supplementary Fig. 8a) and penetrates into the groove of α-β tubulin intradimer (Fig. 4f). Furthermore, α-tubulin H11’-W407, Y408, V409, and G410 form a hydrophobic pocket, into which GAR domain β5 W257 snugly fits into this pocket alongside α- tubulin W407 (Fig. 4g). The density map and atomic models reveal GAR domain α-helix 2 K266 and H267 proximity to β-tubulin E196, suggesting electrostatic interactions (Fig. 4h). Additionally, GAR domain β3 K237 is poised to interact with β-tubulin E196 (Fig. 4h). From the structure analysis, we concluded that the GAR domains interact with microtubules through β3-β4-β5-α2 regions.

The zinc-binding motif is crucial for maintaining the structural integrity of the GAR domain (Fig. 4c, d). We examined the correlation between microtubule (MT) binding and pathogenic GAR mutations documented in the Online Mendelian Inheritance in Man (OMIM) database ^38^. Notably, mutations in the zinc-binding motif of MACF1’s GAR domain are associated with cortical malformations and brainstem anomalies. To simulate the effects of these mutations, we removed the zinc by adding EDTA to the buffer. This zinc depletion significantly diminished the MT binding affinity of the GAS2-GAR protein (Fig. 4i, Supplementary Fig. 3h, i), highlighting the pivotal role of zinc in this interaction.

To validate the GAS2-GAR-MT interactions, we generated a series of point mutants to disrupt observed interactions potentially important for binding. We performed an in vitro co-precipitation assay with taxol-stabilized microtubules (Supplementary Fig. 3h, i). As wild-type GAS2-GAR exhibited *K^d^* of ∼0.69 μM, the GAS2-GAR R242E mutation reduced binding activity to ∼5 μM (Fig. 4i, Supplementary Fig. 3h, i). GAS2-GAR R252E mutation significantly impacted binding to *K^d^* of ∼47 μM (Fig. 4i, Supplementary Fig. 3h, i), suggesting the importance of the interaction between GAS2-GAR R252 and β-tubulin D199. Additionally, mutation W257E resulted in *K^d^* ∼27 μM, suggesting disruption of hydrophobic interactions against α-tubulin W407. Mutating glycine residues at G-hairpin to alanine (G255A) or glutamic acid (G255E) decreased binding ability significantly (*K^d^* ∼2.69 μM for G255A, ∼9.60 μM for G255E, Fig. 4i, Supplementary Fig. 3h, i). GAS2-GAR K266E at α-helix 2 mutation disrupting interfaces with β-tubulin E196 also reduced the binding activity *K^d^* of ∼10.6 μM (Fig. 4i, Supplementary Fig. 3h, i). From these analyses, we concluded that β3-β4-β5-α2 regions are vital for the GAS2 GAR domain to bind to MT.

### GAS2 controls the inter-bundle distance of the MT

In our GAS2-GAR-MT structure, the C-terminal density remained elusive due to its inherent flexibility. We also successfully resolved the full-length GAS2-FL-MT structure at 4.39 Å; however, the C-terminal and the hinge linker between the CH3 domain of the GAS2 protein remained elusive (Supplementary Fig. 4a). This suggests that the flexibility of the hinge linker and C-terminal allows for the optimal orientation of the CH3 and GAR domains when the GAS2 protein mediates the crosslinking of two filaments.

To gain further insights into how GAS2 mediates crosslinking between F-actin and MT, we utilized AlphaFold2 to predict the dimerization structure of the C-terminal (Supplementary Fig. 4b, c) ^36,39^. The final 30 amino acids collectively adopt a plausible dimerization structure (Supplementary Fig. 4b, residues P280 to K314). Notably, the 14 amino acids following the GAR domain (residues R271 to P284) exhibit significant flexibility, potentially contributing to the adjustment of inter-filament distances (Supplementary Fig. 4b). This flexibility prediction aligns with the GAR NMR structure (PDB: 1V5R, Supplementary Fig.3f).

To investigate how GAS2 organizes MT bundles, we compared the polarity of the bundled MT by out scripts (Fig. 5a, b represents the GAS2-GAR domain bundle MT polarity, arrows represent the polarity of MT from minus to plus, Fig. 5d, e represent GAS2-FL). The ratio of parallel and anti- parallel bundle MT pairs has no significant difference (Fig. 5c), which illustrates the ability of the GAS2 protein to bundle MT in either orientation. The bundling MT-to-MT distances observed in the Pillar and Deiters’ cells are about 120-150 Å (Supplementary Fig. 1b) ^40,41^. From our cryo-EM micrographs, the distance between two bundled MTs ranges from 40 to 130 Å in the presence of the GAS2-GAR domain, 70 to 170 Å in the presence of GAS2-FL protein (Fig. 5f). These results indicate that this MT-to-MT distance is consistent with Pillar or Deiters’ cells bundling MT and also indicate the dimerization GAS2 can adjust the inter-filament distances by the flexible loops.

**Fig. 5.**
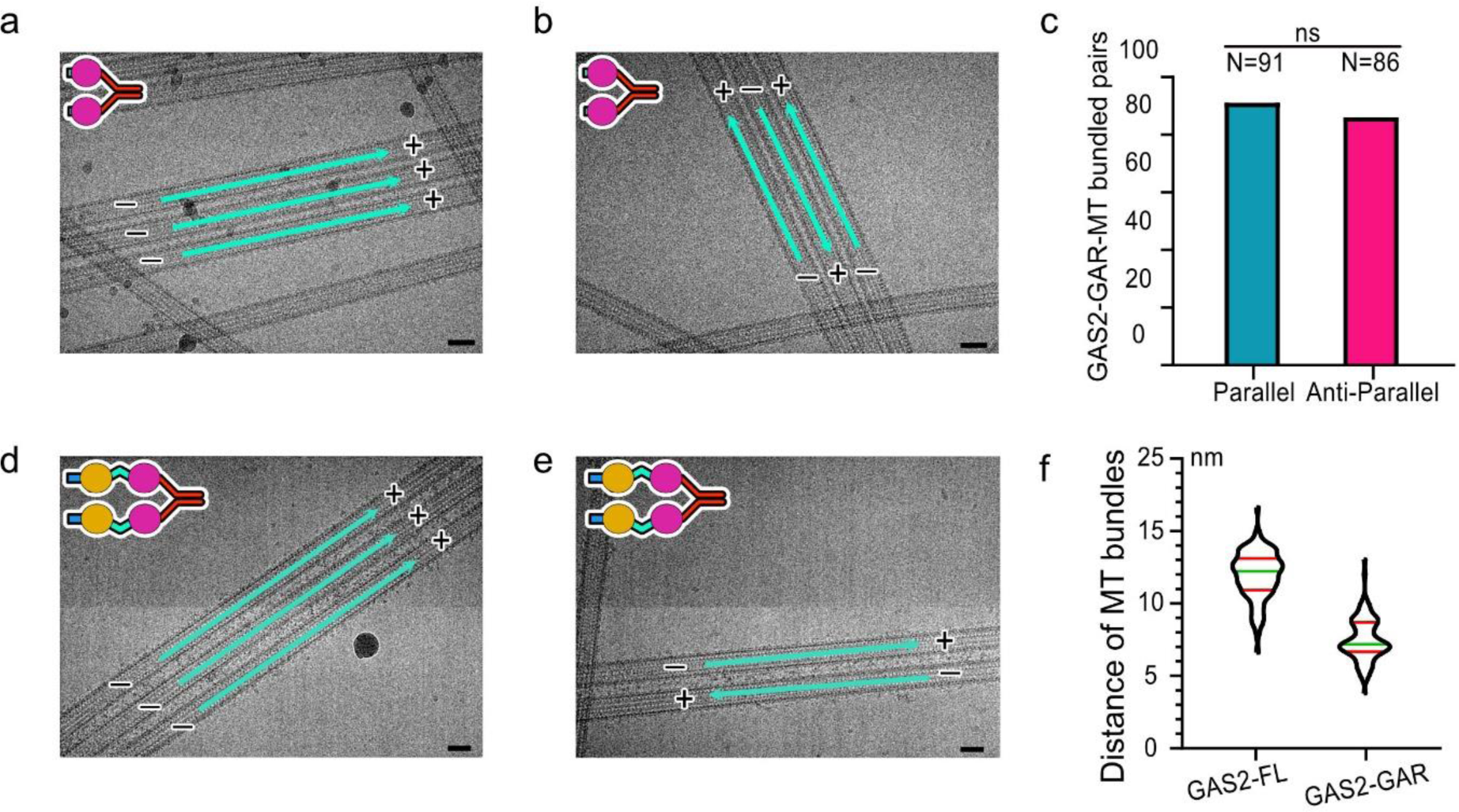
GAS2 controls the inter-bundle distance of the MT. a-b. Microtubule polarity was traced in the GAS2-GAR-MT cryo-EM micrographs, the representative EM images shown the parallel bundled MTs in (**a**), and the antiparallel bundled MTs in (**b**). Cyan arrows represent the polarity of MT from minus to plus. **c** Histogram of GAS2-GAR-MT bundled pairs. Totally, N=177 MT bundling pairs were checked, N=91 MT pairs were bundled in a parallel way, N=86 MT pairs were bundled in an antiparallel way. **d-e** Microtubule polarity was traced in the GAS2-FL-MT cryo-EM micrographs, the representative EM images shown the parallel bundled MTs in (**d**), and the antiparallel bundled MTs in (**e**). Cyan arrows represent the polarity of MT from minus to plus. **f** The distance between the bundled MTs were measured in cryo-EM micrographs. N=63 MT bundled pairs were checked in GAS2- FL-MT cryo-EM micrographs, N=89 MT bundled pairs were checked in GAS2-GAR-MT cryo-EM micrographs. Scale bar = 20 nm. All the detailed values are shown in Source data.

### GAS2-GAR nucleates and stabilizes the MT

We investigated the time-dependent MT formation using negative stain electron microscopy (EM). Initially, 3 μM tubulin was incubated with 3 μM GAS2-GAR for 10 minutes on ice, followed by incubation at 37°C for one minute. This led to the formation of numerous ring structures with diameters of ∼50 nm (Fig. 6a). Over time, we observed a significant increase in microtubules accompanied by a reduction in ring structures (Fig. 6b-d). After 10 minutes at 37°C, the rings disappeared entirely, indicating their utilization in microtubule growth processes (Fig. 6a-d). In contrast, no ring structures were detected in the tubulin-alone control, even after incubation at 37°C for 10 min (Supplementary Fig. 5a, b). Moreover, microtubules were not observed in the tubulin-alone control until 30 minutes of incubation, and even then, only a small number of microtubules were detected (Supplementary Fig. 5c). These results demonstrate that the dimerization of GAS2-GAR induces longitudinal tubulin oligomerization, facilitating microtubule nucleation and polymerization.

**Fig. 6.**
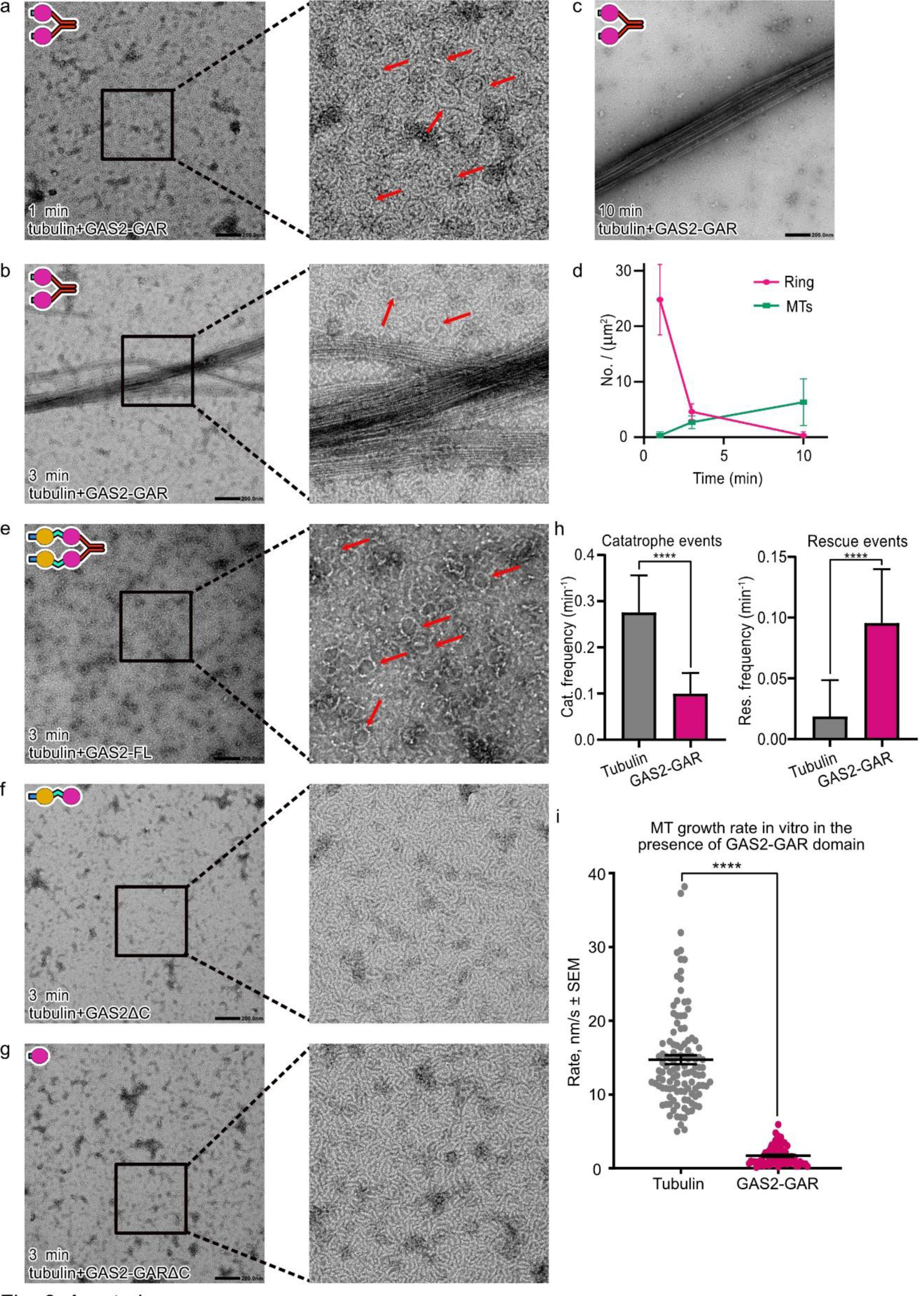
GAS2-GAR nucleates and stabilizes the MT. a-c. Representative negative stain EM micrographs of 3 µM tubulin polymerization with 3 µM GAS2-GAR at different time points. The tubulin rings structure are indicated by red arrows. **d** Plots of the number of tubulin rings (pink) and that of microtubules (green) at different time points (mean ± SD., from 10 independent views). All the detailed values are shown in Source data.**e** Representative negative stain EM micrographs of 3 µM tubulin polymerization with 3 µM GAS2-FL at 3 min. The tubulin rings structure are indicated by red arrows. **f** Representative negative stain EM micrographs of 3 µM tubulin polymerization with 3 µM GAS2ΔC at 3 min. **g** Representative negative stain EM micrographs of 3 µM tubulin polymerization with 3 µM GAS2-GARΔC at 3 min. **h** Effects of GAS2-GAR on frequency of catastrophes and rescues in microtubules (All the detailed values are shown in Source data). Microtubules were assembled from 10 µM tubulin in the presence of 100 nM GAS2-GAR. For control, bare- MT N=19 MTs were checked. For GAS2-GAR N=50 MTs were checked. **** p < 0.0001 (T-tests, nonparametric tests, two-tailed). p values were calculated relative to the tubulin-alone condition. **i** Microtubule growth rate in vitro in the presence or absence of GAS2-GAR- GFP. Black bars show mean ± SEM. For control, bare-MT N=19 MTs were checked. For GAS2-GAR-MT N=52 MTs were checked. **** means t-test p-value<0.0001 compared with control (bare-MT). (T-tests, nonparametric tests, two-tailed). p values were calculated relative to the tubulin-alone condition. All the detailed values are shown in Source data.

Next, to investigate which domains are important, we incubated 3 μM tubulin with 3 μM GAS2- FL, GAS2ΔC, or GAS2-GARΔC. We observed the same ring structures in the presence of GAS2- FL from the negative stain EM micrographs (Fig. 6e). In contrast, no ring structures or microtubules were observed in the presence of GAS2ΔC or GAS2-GARΔC (Fig. 6f, g). These findings suggest that the dimerization of GAS2 constructs has a more prominent effect on tubulin polymerization.

To investigate how GAS2-GAR affects the microtubule dynamics, we conducted total internal reflection fluorescence (TIRF)-based assays with microtubules nucleated from randomly oriented short seeds in the presence of pure tubulin (10 µM) and GAS2-GAR (100 nM) (Supplementary Fig. 5d) and analyzed the frequency of catastrophe (transition from growth to shrinkage) and rescue (transition from shrinkage to growth) events. Compared to tubulin alone, GAS2-GAR significantly inhibited catastrophes and promoted rescue events (Fig. 6h, Supplementary Fig. 5e, f). GAS2-GAR also inhibited the MT growth rate, because microtubules grew at a rate of 14.73±0.60 nm/s in the absence of GAS2-GAR while it is 1.71±0.15 nm/s in the presence of 100 nM GAS2-GAR (Fig. 6i). Interestingly, in the presence of GAS2-GAR, the MTs tend to pause after polymerization, and the average pause time is significantly longer, further illustrating that GAS2-GAR promotes microtubule stabilization (Supplementary Fig. 5g).

### GAS2 crosslinks F-actin and Microtubule

Our structural and biochemical analyses unveil GAS2’s ability to interact with F-actin via the CH3 domain, facilitating filament bundling, and with MT via the GAR domain, promoting MT bundling. This suggests GAS2’s potential as a crosslinker between F-actin and MT. To test this idea, we mixed GAS2, F-actin, and MT, and incubated at 25 °C for 50 minutes, and performed negative stain EM. In the absence of GAS2-FL, there are individual MT and F-actin filaments, but they are not connected nor bundled (Fig. 1l, 3a, 7a). On the other hand, the presence of GAS2-FL led to the formation of F-actin-MT bundles, with large F-actin bundles surrounding the MT (Fig. 7b, d). In the presence of GAS2ΔC, the single F-actin filaments were also tethered to the MT, however, we did not see the F-actin bundling cluster surrounding the MT (Fig. 7c, d).

**Fig. 7.**
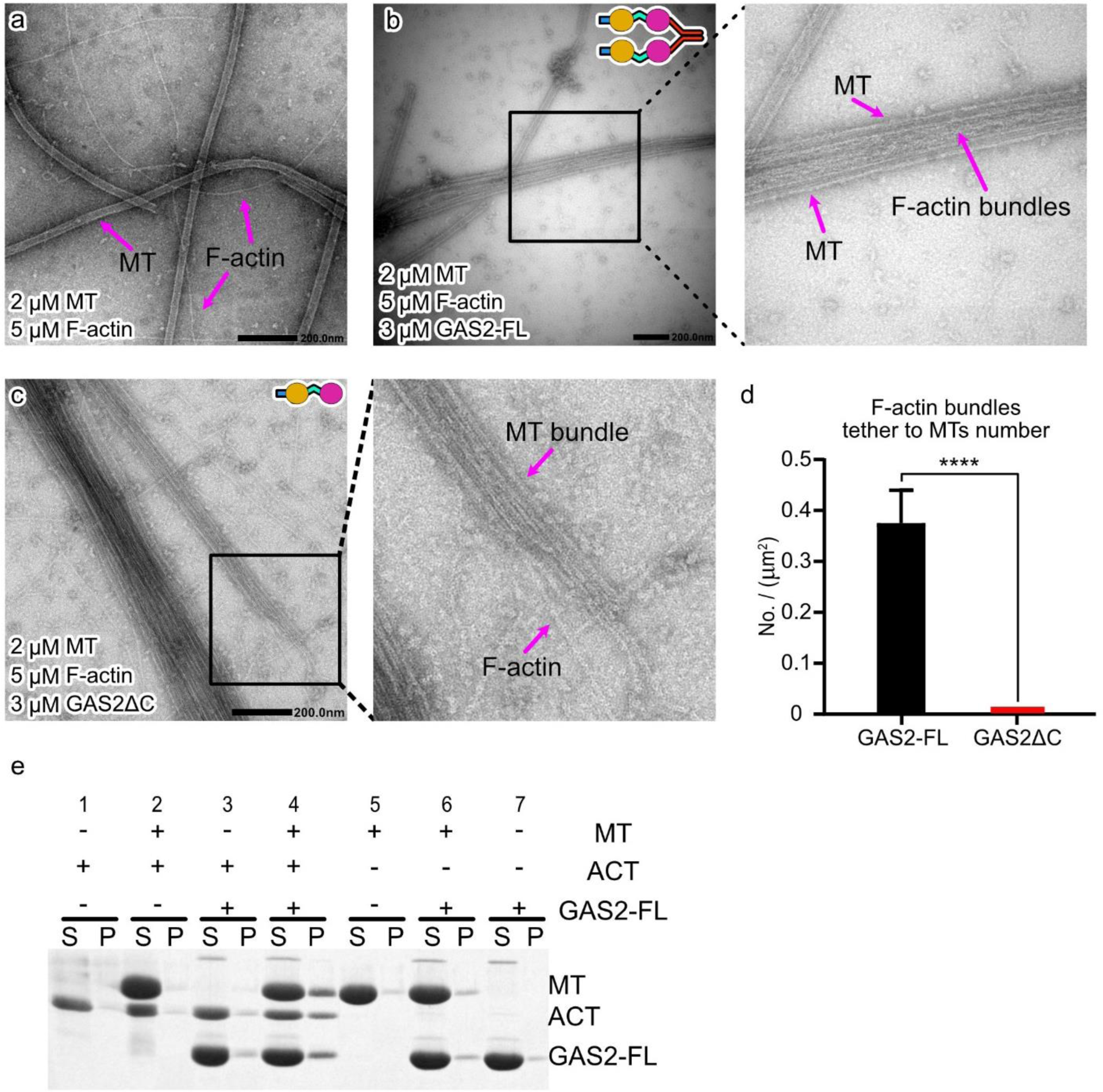
GAS2 mediates F-actin and MT crosslinking. a. Representative negative stain EM micrographs of mixture F-actin and MT are disordered as individual filaments. The MT and F-actin are indicated by arrows. **b** Representative negative stain EM micrographs in the presence of GAS2-FL protein, F-actin, and MT, MT and F-actin bundles were formed, large F-actin bundles surround and along with the MT. The MT and F-actin bundles are indicated by arrows. **c** Representative negative stain EM micrographs in the presence of GAS2ΔC, F-actin, and MTs, F-actin was tethered to MT. The MT and F-actin are indicated by arrows. **d** Histogram of the number of F-actin bundles tethering to MTs (mean ± SEM., from 16 independent views). All the detailed values are shown in Source data. **e** Representative Coomassie-stained low speed co-sedimentation assay gels containing supernatant (S) and pellet (P) for mixture of MT, F-actin and full-length GAS2. Pellet quantitative analysis is shown in (**Supplementary** Fig. 5h-j).

Next, we used low-speed pelleting assays, which can capture only crosslinked actin and MT, to further investigate the crosslinking effects of GAS2 protein on F-actin and MT. After incubating MT and/or F-actin with or without GAS2-FL protein, the mixtures were centrifuged for 10 min at 800 × *g*, diagnosed by SDS-PAGE, and quantified. Small amounts of pelleted actin were observed in F-actin only, MT and F-actin, and F-actin and GAS2-FL reactions (Fig. 7e column 1, 2, and 3, Supplementary Fig. 5h). Similarly, small amounts of tubulin pellet were observed in MT alone, MT and F-actin, and MT and GAS2-FL reactions (Fig. 7e column 2, 5, and 6, Supplementary Fig. 5i). However, in the presence of GAS2-FL, MT, and F-actin, substantial amounts of both actin and tubulin were observed in the pellet (Fig. 7e column 4, Supplementary Fig. 5h-j), suggesting that GAS2 crosslinked actin and MT. These results are consistent with the observation of the GAS2- F-actin-MT complexes by negative staining microscopy.

## Discussion

### GAS2 interacts with F-actin

GAS2, which contains only one CH3 domain, belongs to the 1×CH protein family. Our biochemical analysis revealed that a single CH domain is sufficient for actin binding. Our high-resolution cryo-EM structure of the GAS2-CH3 domain bound to F-actin provides the first example of the single CH3 domain interacting with F-actin.

The structures of CH1 bound to F-actin, such as those in FLNaCH1 (filamin) ^42^, UTRN-ABD (utrophin) ^43^ , and T-plastin ^44^, have been previously elucidated by cryo-EM. A comparison of these CH1 domains with the CH3 domain bound to F-actin reveals significant differences. From the sequence similarity, we found the loops or short helices region of different type CH domains are diverse in length and sequence. First, the CH3 domain tends to have a longer and more well-ordered N-terminal (Supplementary Fig. 6a). The longer N-terminal of the GAS2-CH3 domain forms a ’hook’ structure to catch to the actin (Supplementary Fig. 6b, orange color represents GAS2-CH3 domain). Second, there is a rigid 3^10^ α-helix in the CH3 domain between helix E and F instead of the loops in the CH1 domain (Supplementary Fig. 6a red box). We also found the GAS2-CH3 domain is far away from the actin D-loop because of the rigid 3^10^ α-helix (Supplementary Fig. 6c, orange color represents GAS2-CH3 domain); this way would result in the weaker binding ability for GAS2-CH3 domain with actin. After structure alignment and measurement, the footprint on actin for these CH domains ^45^, we found the T-plastin N-terminal does not interact with actin, FLNaCH1, UTRN-ABD and GAS2-CH3 representatively interact with actin by N-terminal helix and loops (Supplementary Fig. 6b), especially, N-terminal of GAS2-CH3 domain has a larger interaction area with actin (Supplementary Fig. 6g highlight with white circles). While FLNaCH1, UTRN-ABD, and T-plastin have a stronger interaction with actin at ABD2 by the flexible loops, the GAS2-CH3 domain has a rigid 3^10^ α-helix instead of the flexible loops (Supplementary Fig. 6a, c). This decreases the binding area with actin (Supplementary Fig. 6d-g highlighted with a black rectangle).

Our cryo-EM structure shows that the single CH3 domain is sufficient to bind with F-actin. The single EB-type CH domain exhibits a strong binding affinity for the microtubule plus-end and a lower binding affinity for F-actin, demonstrating its selectivity ^46^. Based on the multiple sequences alignment, the EB-type CH domain lacks the ABD2’ binding site, which explains its low affinity for F-actin (Supplementary Fig.7a brown box). The EB-type CH domain also has a shorter loop between helix E and F, which is essential for CH1 and CH3 domains to contact with F-actin (Supplementary Fig.7a red box). These results indicate although the whole globular structure of the CH domain is conserved, the varying loops and short helix empowered the diverse functions of the CH domain.

### GAS2 interacts with MT

The GAR domain is highly conserved from zebrafish to human, especially for the microtubule-binding sites (β3-β4-β5-α2) and zinc-binding motif (Supplementary Fig. 8a). This evolutionarily conserved amino acid sequence predicts the functional importance of this domain. From our GAS2-GAR-MT complex structure, the GAS2-GAR domain specifically binds to the tubulin intradimer interface. Electrostatic analysis revealed complementary charges between the GAR domain and microtubules, with positively charged residues on the GAR domain aligning with negatively charged regions on tubulin intradimer (Supplementary Fig. 8b, c). The GAS2-GAR domain extends deeply into the intradimer concavity, interacting closely with β-tubulin’s loop6- loop8 pocket, particularly aspartic acid 163 (β-D163), while α-tubulin’s lysine 163 (α-K163) prevents binding due to steric hindrance and electrostatic repulsion (Fig. 4e, Supplementary Fig. 8b).

In the presence of GAS2-FL or GAS2-GAR, tubulin rings were produced via the longitudinal growth of αβ-tubulin to form protofilament rings. This mechanism differs from the γ-TuRC, where γ-tubulins contact laterally to form a ring template. Upon incubation at 37°C, these tubulin rings transitioned into sheets and microtubules over time, with a concomitant decrease in ring structures. This observation suggests that GAS2 facilitates tubulin ring formation, with these rings potentially serving as precursors for microtubule formation. However, we did not directly observe the incorporation of tubulin rings into sheets or microtubules. In the presence of GAS2-GAR, we also observed many curved sheets with lower curvatures, indicating that microtubule growth might involve dynamic fluctuations among rings, curved sheets, straight sheets, and microtubules. These results demonstrate that GAS2 can promote tubulin nucleation and microtubule polymerization independently of the γ-TuRC complex and centrosomes.

### GAS2 bundling and crosslinking ability in cell

A previous study demonstrated that Caspase-3 can cleave GAS2 at aspartic residue 279 during cell apoptosis, leading to cytoskeletal rearrangement, changes in cell morphology, and playing a crucial role in chondrogenesis^14,47^. Our fluorescence intensity measurements and negative-stain microscopy reveal that the C-terminal region of GAS2 (residues 277-314) is essential for its F-actin bundling activity. Loss of this C-terminal region abolishes GAS2’s ability to bundle F-actin, resulting in the disruption of F-actin integrity and the transition of F-actin into disordered individual filaments, ultimately causing cytoskeletal rearrangement during cellular processes (Fig. 8a). Additionally, our results show that GAS2 can tether F-actin bundles to microtubules. However, when GAS2 loses the C-terminal region, it loses its F-actin bundling ability. Although it obtains to tether single F-actin filaments to microtubule ability, the crosslinking pattern is altered (Fig. 7b, c). We hypothesize that this deletion imparts a different crosslinking ability to GAS2ΔC, which significantly disrupts cytoskeletal organization, causing cell membrane collapse and cell shape changes (Fig. 8a).

**Fig. 8.**
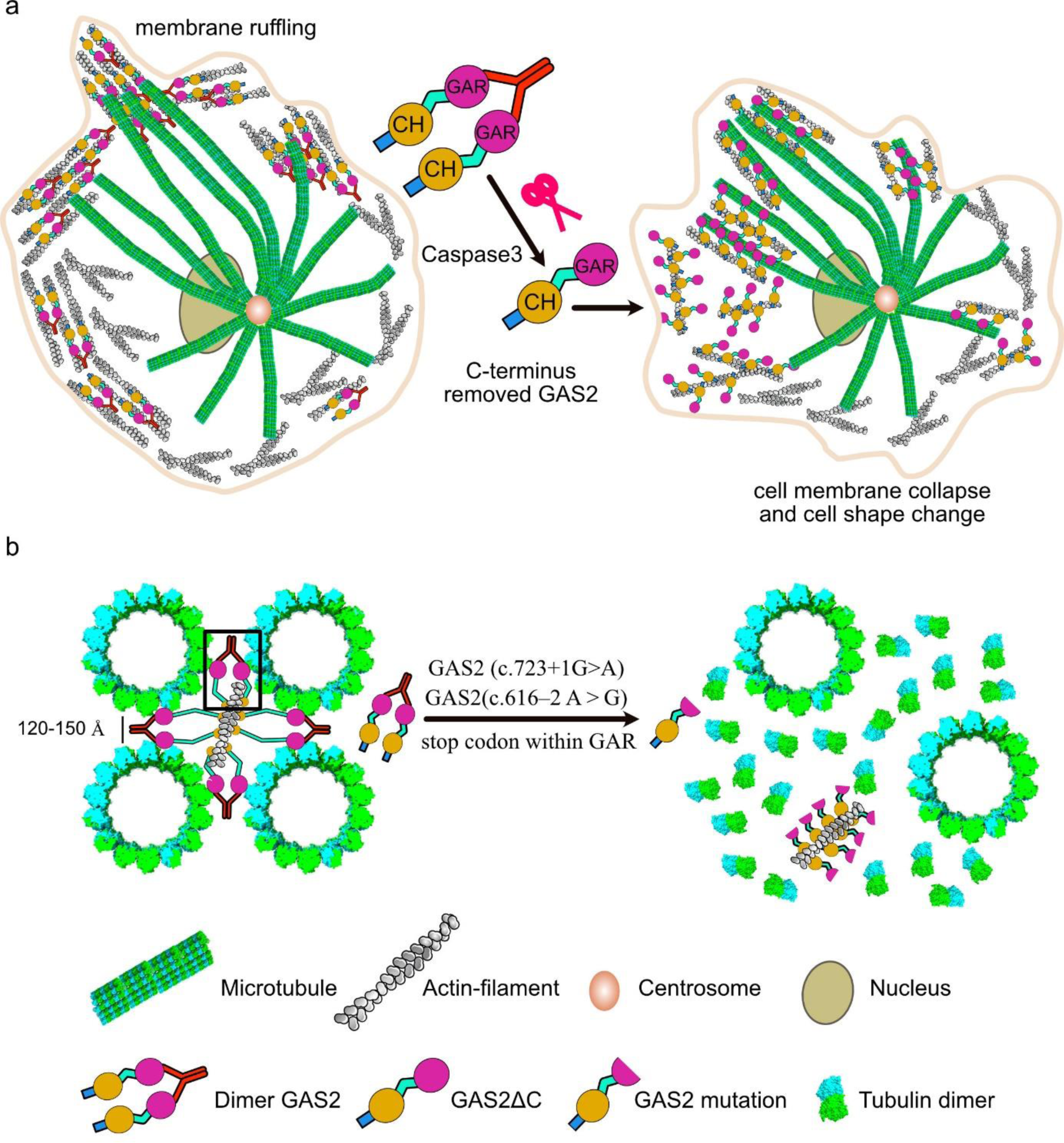
Model of GAS2-driven microtubule bundling or F-actin bundling. a. A simple work model for GAS2-FL in the cells. The GAS2-FL C-terminal being digested by caspase3, removed, or mutated would lead to disordered actin filament arrays or MT-arrays, then causing the specifically differentiated cell shape changing or cell architecture catastrophe. **b** Schematic of GAS2 arranges MT and F-actin in Pillar/Deiters’ cell. Dimer GAS2 is like flexible bridge, four dimer GAS2 bind with MT and link to F-actin form a square. Black box shows one unit dimerization GAS2-FL protein catches two MTs together by two GAR domains, and two CH3 domains tether to the F-actin. The mutations disrupt the GAR domain and delete the C-terminal, causing GAS2 to lose its MT bundling and stabilization ability, resulting in MT depolymerization and MTs arrays catastrophe.

Our structural and biochemical data can explain how GAS2 would work in the inner ear. Pillar and Deiters’ cells are specialized supporting cells with intricate morphologies and cytoskeletal specializations, forming strategic connections with inner and outer hair cells in the organ of Corti (Supplementary Fig. 1b)^48^. In these cells, the microtubules are organized in a square array, interspersed with an equivalent number of actin filaments, with inter-microtubule distances ranging from 120 to 150 Å (Supplementary Fig. 1b)^40^. GAS2 protein colocalizes with both actin filaments and microtubules^33,34^. The dark-field assays and negative-stain electron microscopy indicate that GAS2 can bundle microtubules into arrays in vitro (Fig. 3d-f). Our cryo-EM data demonstrate that, in the presence of GAS2, the distance between microtubules ranges from 70 to 170 Å (Fig. 5d-f), consistent with observations in Pillar cells, suggesting a role for GAS2 in organizing microtubule distances. GAS2 may act as a flexible bridge in Pillar cells, eight GAR domains of four dimerization GAS2 pairs bundle four MTs at an optimal distance to form a square, and the CH3 domains tethering to F-actin, thus a perfect MTs-F-actin crosslinking arrays are formed (Fig. 8b, black box shows one of the GAS2-FL dimer units, Supplementary Fig. 1b).

Our biochemical data can explain the genetic diseases caused by GAS2 mutations. Variants such as GAS2 c.723+1G>A and GAS2 c.616–2A>G introduce a stop codon within the GAR domain, resulting in disordered microtubule arrays, microtubule depolymerization, and inherited hearing loss^33,34^. Our TIRF data shows that GAS2-GAR reduces catastrophe and promotes the rescue of MTs, stabilizes MTs. However, the variants disrupting the GAR domain would impair GAS2’s ability to stabilize MTs, leading to microtubule depolymerization. However, microtubule bundle arrays are not formed when GAS2 lacks the C-terminal region (Fig. 3b), indicating a loss of MT bundling ability in Pillar and Deiters’ cells. Variants such as GAS2 c.723+1G>A and GAS2 c.616– 2A>G lead to C-terminal deletion and GAR domain disruption, causing GAS2 to lose its MT bundling ability, impairing stabilization, and resulting in MT depolymerization and MTs arrays catastrophe (Fig. 8b). Additionally, there are three non-centrosomal microtubule-organizing centers (nc-MTOCs) in Pillar cells generating MT bundle arrays (Supplementary Fig. 1b) ^49^. The GAS2 C-terminal region is crucial for ring structure formation. The truncation of this region results in the disappearance of ring structures and reduced microtubule polymerization compared to GAS2-FL or GAS2-GAR constructs. These results indicate that GAS2-FL potentially assists in nc-MTOC formation, and C-terminal deletion may also impair this function.

In summary, our research revealed that GAS2 protein can promote MTs nucleation, facilitate MTs stabilization, bundle MTs or F-actin together, and crosslink F-actin bundles to MTs. Our research significantly enhances the molecular diagnosis of hearing loss and lays the groundwork for future studies to develop new therapeutic strategies to prevent progressive hearing loss linked to GAS2.

## Materials and Methods

### Cloning, Expression, and Recombinant Protein Purification

Mouse GAS2 constructs were subcloned into modified pCold-Pros2 vectors, where the Thrombin and Factor Xa sites had been removed. GAS2-FL (1-314), GAS2ΔC (1-276), GAS2-CH3 (1-201), GAS2-CH3ΔN (34-201), GAS2-GAR (198-314), GAS2-GARΔC (198-276) and their mutants were inserted into pCold-ProS2 after an HRV3C cleavage site (LGVLFG/GP). For GFP-tagged constructs, EGFP was inserted at the C-terminus of the protein within the pCold-ProS2 plasmid. Proteins were expressed in BL21(DE3) at 20°C for 16 h following induction with 0.5 mM IPTG at OD600 = 0.6. Bacteria were lysed by sonication in buffer A (30 mM HEPES, pH 7.5, 300 mM KCl). Lysates were centrifuged at 20,000 × g for 30 min at 4°C. Ni-NTA chromatography was used for protein purification. ProS2 tags were removed by on-column digestion with HRV3C protease in buffer A. GAS2-FL, GAS2ΔC, GAS2-GAR, GAS2-GARΔC, and their mutants were dialyzed against buffer B (25 mM Tris-HCl, pH 8.0, 150 mM KCl, 100 μM ZnCl^2^, 1 mM DTT). GAS2-CH3, GAS2-CH3ΔN, and their mutants were dialyzed against Tris-Triton buffer (50 mM Tris-HCl, pH 8.0, 200 mM KCl, 2 mM MgCl^2^, 1 mM DTT, 0.01% Triton X-100). GFP-tagged constructs were purified using the same protocol as their non-GFP counterparts.

### Actin and Tubulin Preparation

Rabbit skeletal muscle actin was prepared from acetone powder (gift from Hirokawa lab) as described previously (Spudich and Watt, 1971). Briefly, actin was extracted from the acetone powder, polymerized with 0.1 M KCl, and then further purified with high-molar KCl (0.6 M) and depolymerization in the low salt buffer. F-actin was polymerized in F-buffer (50 mM Tris-HCl, pH 8.0, 200 mM KCl, 2 mM MgCl^2^, 1 mM EGTA, 4 mM DTT). Tubulin was purified from porcine brain using a high-molarity PIPES buffer (Castoldi and Popov, 2003). GMPCPP-stabilized microtubules were prepared by polymerizing tubulin with GMPCPP as described by Yajima et al. (2012) ^50^. Taxol-stabilized microtubules were prepared by polymerizing 5 mg/mL tubulin in the BRB-80 buffer (80 mM PIPES, pH 6.8, 2 mM MgCl^2^, 1 mM EGTA, 1mM DTT) with 50 mM GTP at 37°C for 30 min, followed by the gradual addition of Taxol to a final concentration of 20 μM.

### Size exclusion chromatography (SEC) assays

SEC assays were performed using a Superdex 75 H.R. 10/300 column on an AKTA purifier system. GAS2-FL, GAS2ΔC, GAS2-GAR, and GAS2-GARΔC were loaded with SEC buffer E (30 mM Tris-HCl, pH 8.0, 300 mM KCl, 100 μM ZnCl^2^, 0.1% β-ME). Eluted fractions were analyzed by SDS-PAGE. Standard proteins (BSA, 66 kDa; Carbonic Anhydrase, 29 kDa; Cytochrome c, 12.4 kDa) were used to generate a calibration curve, which was then used to estimate the molecular weights of GAS2-FL and its constructs based on their elution volumes (Supplementary Fig. 1c-e).

### Actin Binding and Bundling Assays

High-speed co-sedimentation assays were performed to investigate the interaction between GAS2 constructs and F-actin. Various concentrations of GAS2 proteins were incubated with pre-polymerized F-actin (5 μM) in the F-buffer for 1 hour at room temperature. The mixtures were then centrifuged at 186,000 × g for 10 min at 25°C. Supernatants and pellets were separated and analyzed by SDS-PAGE. The amount of GAS2 proteins in the pellet fractions was quantified using ImageJ software (NIH) to determine the binding affinity.

Low-speed co-sedimentation assays were conducted to assess the F-actin bundling activity of GAS2 constructs. F-actin (5 μM) was incubated with various concentrations of GAS2 proteins in the F-buffer for 1 hour at room temperature. The mixtures were centrifuged at 9,000 × g for 10 min at 4°C. Supernatants and pellets were separated and analyzed by SDS-PAGE. The percentage of actin in the pellet fractions was quantified to evaluate the bundling efficiency of each construct.

The F-actin bundles induced by GAS2 were visualized using fluorescence microscopy and negative stain electron microscopy. For fluorescence microscopy, F-actin (5 μM) was incubated with GAS2-FL (1, 3, and 5 μM) in F-buffer for 1 hour at room temperature. The mixture was diluted 10-fold with imaging buffer (50 mM Tris-HCl, pH 8.0, 200 mM KCl, 2 mM MgCl^2^, 1 mM EGTA, 4 mM DTT, 50 μL/mL Alexa Fluor 568 phalloidin), applied to poly-L-lysine coated coverslips, and imaged using a FV3000 (Olympus Life Science) confocal microscope. For negative stain electron microscopy, the same procedure was followed as for fluorescence microscopy, except that the samples mixture was diluted with F-buffer and applied to glow-discharged carbon-coated copper grids, stained with 1% uranyl acetate, and imaged using a JEOL JEM-1400 transmission electron microscope operated at 100 kV.

### Microtubule Binding and Bundling Assays

Co-pelleting assays were performed to confirm the binding affinity of GAS2-FL and its GAS2- GAR domain (198-314) to microtubules. Different concentrations of GAS2 proteins were incubated with Taxol-stabilized MTs in BRB80 buffer supplemented with 1 mM GTP at room temperature for 20 min. The mixtures were then centrifuged at 186,000 × g for 10 min at 25°C. All experiments were performed in triplicate, then supernatants and pellets were separated and analyzed by SDS-PAGE. The gels were quantified using ImageJ software (NIH), background subtraction was performed using the respective GAS2 construct controls, and statistical analysis was carried out using GraphPad (GraphPad Software, San Diego, USA).

To examine the effect of Zn^2+^ on the GAS2-GAR domain’s interaction with microtubules, co-sedimentation assays were performed under two conditions: The GAS2-GAR domain was dialyzed into either buffer D (25 mM Tris-HCl, pH 8.0, 100 mM KCl, 100 μM ZnCl^2^, 1 mM DTT) for the “With Zn^2+^” condition or buffer-EDTA, which has the same composition as buffer D except that ZnCl^2^ was replaced with 10 mM EDTA, for the “Without Zn^2+^” condition. The co-sedimentation assays were performed and analyzed as described above.

Dark-field microscopy and electron microscopy were used to observe MT bundling induced by GAS2 proteins. Taxol-stabilized MTs (2 μM) were incubated with 1.0 μM GAS2 proteins (GAS2- FL, GAS2ΔC, GAS2-GAR, GAS2-GARΔC) in BRB80 at room temperature for 30 min. The samples were then imaged using either dark-field microscopy (BX53, Olympus) or negative stain electron microscopy as described for the F-actin bundling assay above.

### Full-length GAS2/F-actin/MT cross-linking assay

Taxol-stabilized MTs and F-actin were prepared as described in the “Actin and Tubulin Preparation” section. 5 μM taxol-stabilized MTs, 5 μM F-actin, and 25 μM GAS2 proteins were mixed in BRB80 and incubated for 40 min at room temperature. After centrifugation at 800 × g for 10 min, supernatants (S) and pellets (P) were separated and analyzed by SDS-PAGE. Control experiments included 5 μM bare MTs, 5 μM bare F-actin, 5 μM bare MTs mixed with 5 μM bare F-actin, 5 μM MTs mixed with 25 μM full-length GAS2, 5 μM F-actin mixed with 25 μM full-length GAS2, and 25 μM full-length GAS2 alone.

For negative stain electron microscopy to examine the F-actin/MT cross-linking ability of full-length GAS2, 2 μM MTs and 5 μM polymerized actin filaments were incubated with 5 μM full-length GAS2 in BRB80 for 40 min at room temperature. The mixture was then diluted by adding 5 μL of the mixture to 45 μL BRB80. Negative staining and imaging with electron microscopy were performed as described for the F-actin bundling assay above.

### Cryo-EM Sample Preparation and Data Collection

For F-actin-GAS2-CH3 complex, 40 μL of Actin (4 μM polymerized in F-buffer) was mixed with 20 μL of 56 μM GAS2-CH3 at room temperature for 40 minutes in F-buffer, followed by the addition of another 20 μL of 56 μM GAS2-CH3 for further incubation for 40 minutes. 3 μL of the mixture was applied to a glow-discharged carbon-coated grid (Quantifoil R1.2/1.3 Au300 mesh) and incubated for 60 s in the chamber of a Vitrobot (22°C, 100% relative humidity; Thermo Fisher Scientific). The grid was blotted for 4 s at force 10 and plunged into ethane immediately.

For MT-GAS2-GAR complex, 4 μL of 0.5 μM GMPCPP-microtubules were applied to a grid and incubated for 30 s in the chamber of a Vitrobot. Then 2 μL of 5 μM GAS2-GAR in BRB80 were added on the grid and incubated for 30 s to allow GAS2-GAR binding to microtubule. 4 μL of the protein mixture was removed from the grid, and another 2 μL of GAS2-GAR was added. After another 30 s incubation, the grid was blotted for 3 s at force 15 and plunged into ethane immediately.

Images were recorded at 300 keV using a Titan Krios G3i microscope (Thermo Fisher Scientific). Images were collected with a Gatan-LS Energy Filter (Gatan, Pleasanton, CA) with a slit width of 20 eV at a magnification of 105,000× and a physical pixel size of 0.83 Å/pixel for F-actin-GAS2- CH3 complex or 0.85 Å/pixel for MT-GAS2-GAR. For F-actin-GAS2-CH3 complex, each movie was recorded with a total dose of 48.93 electrons/Å2 and subdivided into 49 frames. Movies were acquired using EPU software (Thermo Fisher Scientific), and the target defocus range was set from -2.4 to -1.2 μm. A total of 6,378 micrographs were collected. For MT-GAS2GAR complex, images were collected in the same condition as above. Total dose of 48.79 electrons/Å2 and subdivided into 48 frames. Movies were acquired using SerialEM software ^51^, and the target defocus range was set from -1.8 to -1 μm. A total of 3,322 micrographs were collected.

### Image Processing and 3D Reconstruction

For both datasets, beam-induced motion correction was performed using MotionCor2 ^52^ with 3 × 3 patches, and the CTF was estimated using CTFFIND4 ^53^.

For F-actin-GAS2-CH3 complex, 5,756 micrographs were selected by subset selection in RELION ^54^ with a maximum metadata value of 5.5. In total, 29,620 segments were manually picked to avoid selecting F-actin filaments without GAS2-CH3 binding. 632,290 particles were extracted with a 27.4 Å helical rise. Several cycles of 2D classification and 3D classification were performed, and a final set of 412,176 particles was used for 3D model reconstruction. After polishing, the final resolution achieved was 2.8 Å (Supplementary Fig. 2a).

For MT-GAS2GAR complex, PyfilamentPicker scripts were used to select 14-PF MTs and cut each MT into overlapping boxes with an 82-Å rise ^55,56^. All selected MTs were stacked into “super-particles”. FREALIGN version 9 was used to determine the helical parameters (twist and rise) for the three-start helix of microtubules from the C1 reconstruction ^57,58^. Based on “super-particles” and seam search scripts, the seam location for each particle was determined after several cycles of FREALIGN local/global refinements. During processing, poorly decorated microtubules and particles with incorrect seam locations were removed. After iterative refinement, the seam location was visible in the C1 reconstruction. From the FREALIGN estimation, the resolution of the density map was 4.43 Å by 0.143 FSC value. Based on this map, we fitted the atomic structure of the tubulin dimer and NMR structure of the GAS2-GAR (PDB: 1V5R) domain into the map and produced an initial atomic model. we copied this atomic model and filled all these models into the 3D density map, and then created 5 Å reference maps (Supplementary Fig. 3c) by Chimera (UCSF, https://www.cgl.ucsf.edu/chimera/) for the subsequent step analysis.

To further improve the resolution, the data was processed following the MiRP method based on the RELION pipeline ^37^ (Supplementary Fig. S3c). We used the same data set and manually picked MT filaments by RELION v3.1, and then 4× binned particles were extracted. MT particles were assigned to 11-16 protofilament microtubules by 3D classification in RELION, and 14 protofilament MTs occupying 90% of the total picked MT were chosen for further reconstruction. Reference maps rebuilt from the FREALIGN refinement described above were used for initial seam alignment and seam checking. The final resolution and the B-factor for map sharpening was estimated using relion_postprocess in RELION-3.1 (Table 1). Due to the presence of the seam, the MT reconstruction contains only one “good” protofilament at the opposite seam position ^37,55^, which was chosen for atomic model building.

### Model Building and Refinement

For F-actin-GAS2-CH3 complex, an atomic model was constructed using the AlphaFold2 ^36^ prediction of the GAS2-CH3 domain (AF2 model No. AF-P11862-F1) as a starting model fitted to density map by Chimera X (https://www.rbvi.ucsf.edu/chimerax/). The majority of the GAS2- CH3 domain fit well into the density map, except for a few flexible loops. Initial atomic coordinates were generated and refined via molecular dynamics flexible fitting (MDFF) ^59,60^ to optimize the fit to the cryo-EM density map. The detailed procedure for MDFF is described below. Manual adjustments were made in Coot to improve the fit. Further refinements were performed using the real-space refinement tool in Phenix ^61^. The atomic model was validated using the validation and map-based comparison tools in Phenix. The statistics of the final atomic models are summarized in Table 1. All the figures were prepared by PyMol (https://pymol.org/), Chimera or ChimeraX.

For MT-GAS2-GAR complex, a single unit of tubulin dimer/GAR complex was extracted from the original density map. The NMR structure of the GAS2-GAR domain (PDB: 1V5R) and tubulin dimer structure (PDB: 3JAT) were rigidly fitted into the density map using Chimera X. Initial atomic coordinates were generated same as above. Given the limited resolution and density in certain GAR domain loops, model building focused on the interface amino acids with tubulin. Further refinements and validation are the same as above.

### Molecular dynamics simulation (MDFF)

The rigid fitted initial atomic model (F-actin-GAS2-CH3 or GAS2-GAR-MT) was loaded into VMD software to prepare the PSF file. Then added a solvation box for the model. Following the next step, the model was ionized with neutralized NaCl ions. After that, the model was performed MDFF by using NAMD software, 50,000 steps were used with 0.3 kcal/mol force-scaling factor for cryo-EM density map and with the presence of secondary structure restraints and cis-peptide restraints. Then the model was performed the energy minimization for 2,000 steps using a force-scaling factor of 10.0 kcal/mol.

### Microtubule polarity trace

Based on the psi and tilt angle of the particles after rebuilding the 3D GAS2-GAR-MT structure, we developed a script to trace the MTs polarity in raw cryo-EM micrographs. Upon executing the script, a directional arrow emerges along the microtubule. The orientation of this arrow in the micrographs corresponds to a three-dimensional vector in Chimera, representing the direction from the negative to the positive along the Z-axis.

### Time-dependent of microtubule nucleation assay by Electron Microscopy

Time-dependent of microtubule nucleation assay by Electron Microscopy as described previously ^62,63^. Briefly, 3 μM tubulin was incubated with 3 μM GA2-FL, or GAS2-GAR, or GAS-FLΔC, or GAS2-GARΔC for 10 minutes on ice in BRB80 buffer. Then the mixture was transferred to 37°C for 1, 3, 10, or 30 min. After incubation, 3 µl of each sample was applied to glow-discharged carbon-coated copper grids, stained with 1% uranyl acetate, and imaged using a JEOL JEM-1400 transmission electron microscope operated at 100 kV.

### MT dynamic assay

The in vitro MT dynamics was observed under the TIRF microscope as described previously^64^. The flow chamber was assembled using microscopy slides and precleaned glass coverslips with double-sided tape. The chamber was treated with 0.5 mg/mL PLL-PEG-biotin in BRB80 buffer for 5 min, washed with BRB80 buffer, and then incubated with 0.5 mg/mL streptavidin for 5 min. Short MT seeds were prepared using a 1.5 μM tubulin mix containing 50% biotin-tubulin and 50% AZdye647-tubulin with 1 mM GMPCPP at 37°C for 30 min. The MT seeds were attached to the coverslips using biotin-avidin links and incubated with assay buffer.

The reaction mixture containing a specified amount of GAS2-GAR-GFP (100 nM), 10 μM tubulin, 1 mM GTP, an oxygen scavenging system composed of Trolox/PCD/PCA, and 2 mM ATP was added to the flow chamber. The samples were maintained at 30°C during the experiments. An ECLIPSE Ti2-E microscope equipped with a CFI Apochromat TIRF 100XC Oil objective lens, an Andor iXion Life 897 camera, and a Ti2-LAPP illumination system (Nikon, Tokyo, Japan) was used to observe single-molecule motility. NIS-Elements AR software ver. 5.2 (Nikon) was used to collect movie data for 20 min.

## Supporting information

Description of Additional Video Files

## Data availability

Data supporting the findings of this paper will be available from the corresponding authors upon reasonable request.

The source data underlying Figs. 2k, 3j, 4i, 5c, 5f, 6d, 6h, 6i, 7d, Supplementary Fig. 1f-j, 2c-d, 3h-i, 5g-j are provided as a Source Data file.

The data generated in this study are available in the following databases: PDB-XXXX (GAS2-CH3-F-actin atomic model)

PDB-XXXX (GAS2-GAR-MT atomic model) EMD-XXXXX (GAS2-CH3-F-actin 3D volume) EMD-XXXXX (GAS2-GAR-MT 3D volume) EMD-XXXXX (GAS2-FL-MT 3D volume)

### Acknowledgements

This work was supported by JSPS KAKENHI Grant Number 21H04762 and 21H05248 to M.K., 21H05254 to R.N., 22H05523 to S.N., and 23K27145 to A.N.. This work was supported by the Platform Project for Supporting Drug Discovery and Life Science Research (Basis for Supporting Innovative Drug Discovery and Life Science Research (BINDS)) from AMED under Grant Number JP22ama121002j002 (to M.K.).

Supplementary Fig. 1b was drawn using images from Servier Medical Art. Servier Medical Art by Servier is licensed under a CC BY 4.0 Unported License (https://creativecommons.org/licenses/by/4.0/).

## Author contributions

Author contributions J.A. and M.K. conceived the project. J.A. carried out the protein production and chemistry experiments. J.A., T.I., R.N., and T.M. performed the GAS2-GAR-MT cryo-EM analyses. J.A. and A.N. performed the GAS2-CH3-F-acrtin cryo-EM analyses. J.A. and S.N. performed the TIRF assay. J.A. and R.S. performed MT polarity analysis. All authors discussed the data. J.A., T.M., and M.K. wrote the paper.

