## Supplementary Figure legend for "Dimerization GAS2 mediates microtubule and F-actin crosslinking"

**Supplementary Fig. 1 a** Domain structure of spectraplakins. **b** The schematic diagram illustrates the inner ear. Sound waves entering the outer ear vibrate the eardrum, with these vibrations amplified by the ossicles (malleus, incus, and stapes) and transferred to the oval window. The oval window transmits the vibrations to the cochlea, where fluid movement creates waves that cause the basilar membrane to vibrate (left panel). These vibrations are transduced into inner and outer hair cells (IHC, OHC) to generate an electrical signal, supported by Pillar and Deiters' cells (middle panel) [1]. Pillar and Deiters' cells feature a robust cytoskeleton with microtubules in tightly bundled square arrays spaced 12-15 nm apart and crosslinked with F-actin (left panel), this intricate structure is vital for sound wave oscillation[2–4]. Three ncMTOCs (non-Centrosomal Microtubule-Organizing Centers) collaborate to coordinate control of microtubule positioning in the Pillar cells (middle panel)[5]. **c** SEC (Size Exclusion Chromatography) results for GAS2-FL, solid red line; GAS2ΔC, solid purple line; GAS2-GAR domain, solid green line; and GAS2-GARΔC, solid blue line with marker protein (black dot line). **d** The calibration formula for Superdex 75 HR 10/300 column according to the peak location of the marker proteins. **e** Calibration standard M.W. line for Superdex 75 HR 10/300 column according to peak location of marker proteins, shown as black dots and pink line. The blue dot lines show the relation between the peak location and the molecular weight derived from the calibration standard line. **f-g** Representative Coomassie Blue-stained gels of co-sedimentation assays, containing supernatant (S) and pellet (P) for purified GAS2-FL and GAS2-CH3 domain. **h-i** Low-speed co-sedimentation assay of purified full-length GAS2 (GAS2-FL) and C-terminal truncation GAS2(1-276). Various amounts of full-length GAS2 or GAS2 (1-276) were mixed with F-actin, incubated for 60 min at room temperature, and then centrifuged for 10 min at 9,000 × g. Both supernatant (S) and pellet (P) were separated on SDS/PAGE and stained with CBB. **j** Co-sedimentation assay for GAS2-CH3 domain with MT and GAS2-GAR domain with F-actin. The GAS2-CH3 domain did not co-pellet with MT, and the GAS2-GAR domain did not co-pellet with F-actin. All the raw SDS-PAGE gels are shown in source data.

**Supplementary Fig. 2. a** F-actin-GAS2-CH3 structure reconstruction workflow by RELION v4.0. **b** Left most panel shows the atomic models for the GAS2-CH3 domain fitted into the

3D reconstruction density map. Actin subunits (marked as n series from bottom to top) are presented in green, indian red, medium orchid, royal blue, and GAS2-CH3 domain is presented in orange, respectively. The right panels show examples of atomic models from subunits built into the experimental cryo-EM maps. **c** The $K_d$ value for WT and mutants of the GAS2-CH3 domain according to co-pellet assay. Each $K_d$ value represents the mean ± SD. from three independent experiments. **d** The fitting curve for WT and mutants of the GAS2-CH3 domain for co-pellet assay. One site-specific binding method was used for fitting the curve. Each data point represents the mean ± SD. from three independent experiments. All the raw SDS-PAGE gels are shown in source data.

**Supplementary Fig. 3. a-b** Representative Coomassie-stained MT co-sedimentation assay gels containing supernatant (S) and pellet (P) for purified GAS2-FL and GAS2-GAR domain. **c** MT-GAS2-GAR structure rebuilding workflow. **d** The 2D class average for a GAS2-GAR decorated on a microtubule. GAS2-GAR domains are regularly spaced at 8 nm. **e** Left panel displayed local resolution estimation of the asymmetric units by RELION v3.1.3. The right panel shows examples of atomic models from subunits built in the experimental cryo-EM maps. **f** One represents NMR structure of GAS2-GAR domain (PDB: 1V5R, residues A199 to P284). **g** Outside top view of atomic models for GAS2-GAR domain are fitted in 3D reconstruction density map (α-tubulin in green, β-tubulin in cyan, GAS2-GAR domain in hot-pink). **h** The Kd value for WT and mutants of the GAS2-GAR domain according to co-pellet assay. Each Kd value represents the mean ± SD from three independent experiments. **i** The fitting curve for WT and mutants of the GAS2-GAR domain for co-pellet assay. One site-specific binding method was used for fitting the curve. Each data point represents the mean ± SD from three independent experiments. All the raw SDS-PAGE gels are shown in source data.

**Supplementary Fig. 4. a** 4.4 Å cryo-EM structure of one microtubule decorated with GAS2-FL. **b** Alpha-fold2-predicted C-terminal region, showing the dimer structure, colored by rainbow from N- to C-terminal. The region from R271 to P284 consists of flexible loops, while the region from P284 to K314 is the dimerization area. **c** The PLDDT score of the rank1 structure in (**b**).

**Supplementary Fig. 5. a-c** Representative negative stain EM micrographs of 3 μM tubulin-alone polymerization at different time points. **d** Schematic of TIRF microscopy-based MT stabilization assay. The biotin-labeled microtubule seed (cyan) was tethered to the cover glass by streptavidin and PLL-PEG biotin. The free tubulin dimers (shown in red and orange color) polymerize or depolymerize at the end of the microtubule seed. **e** Representative kymographs of microtubules (magenta) growing from GMPCPP seeds (cyan) in tubulin-alone, black arrow shows one of the catastrophe events. Horizontal scale bars: 5 μm; vertical scale bars: 1 min. **f** Representative kymographs of one microtubule (magenta) growing from GMPCPP seeds (cyan) in the presence of GAS2-GAR-GFP (green). Yellow stars represent the rescue event. From left to right, the panels show the seeding, GAS2-GAR-GFP, tubulin (magenta), and the merged kymographs. Horizontal scale bar: 5 μm; vertical scale bar: 1 min. **g** Left panel shows the representative kymograph of microtubules pause state in the presence of GAS2-GAR. The yellow stars represent the rescue event. Growth represents MT polymerization. The right panel shows the average pause time per microtubule. Black bars show mean ± SEM. For control, bare-MT N=19 MTs were analyzed. For GAS2-GAR, N=50 MTs were analyzed. **** $p < 0.0001$ (T-tests, nonparametric tests, two-tailed). p values were calculated relative to the tubulin-alone condition. Horizontal scale bar: 5 μm; vertical scale bar: 1 min. **h** Histogram of pellet quantitative analysis for F-actin in (**Fig. 7e**). **i** Histogram of pellet quantitative analysis for MT in (**Fig. 7e**). **j** Histogram of pellet quantitative analysis for full-length GAS2 in (**Fig. 7e)**. Each value represents the mean ± SD. from three independent experiments. One-way ANOVA with Dunnett's multiple comparisons test has been used. **, p-value < 0.01, ***, p-value < 0.001.

**Supplementary Fig. 6. a** CH1 and CH3 domain sequence alignment. Primary accession on Uniprot for each sequence: GAS2-CH3 (mouse): P11862; Dystrophin (human): P11532; Calponin-1 (human): P51911; Utrophin (human): P46939; ACTN1 (human): P12814; ACTN2 (human): P35609; SPTBN2 (human): O15020; FilaminA (human): P21333. (F) CH3 domain sequence alignment. Primary accession on Uniprot for each sequence: GAS2CH (mouse): P11862; mal3 (yeast): Q10113; EB1 (mouse): Q61166; EB2 (mouse): Q8R001; EB3(mouse): Q6PER3; Calponin-1(human): P51911; Tagln2 (mouse) Q9WVA4. ABD1 sequences were shown in the green box. ABD2' sequences were shown in the brown box. $3_{10}$ α-helix was shown in the red box. ABD2 sequences were shown in the black box. **b** CH domain N-terminal structure alignment, GAS2-CH3 (orange) interacts with actin by a longer N-terminal compared with T-plastin (silver, PDB:7R94, EMDB: EMD-24323) [6], utrophin (gray, EMDB: EMD-30085, PDB: 6M5G) [7] and FLNA (wheat, EMDB: EMD-7831, PDB:6D8C) [8]. **c** GAS2-CH3 domain rigid $3_{10}$ α-helix structure compares with the loops of T-plastin, utrophin, and FLNA by the same color scheme and structure data bank source as in (**b**). **d-g** Footprint method to inspect potential contacts between actin and the binding partner as defined within 5-Å distance by ChimeraX with the same structure data bank source as in (**b**).

**Supplementary Fig. 7 a** F-actin binding single CH3 domain and MT binding sequence alignment. Primary accession on Uniprot for each sequence: GAS2-CH3 (mouse): P11862; Calponin-1 (human): P51911; Tagln2 (mouse): Q9WVA4, Mal3 (yeast): Q10113; EB1 (mouse): Q61166; EB2 (mouse): Q8R001; EB3 (mouse): Q6PER3; EB3 (human): Q9UPY8. ABD2' sequences were shown in the brown box. Loop sequences between E and F helix were shown in the red box.

**Supplementary Fig. 8. a** Sequence alignment of the GAR domain from human, mouse, Drosophila melanogaster, *C. elegans,* and zebrafish. The black line shows the conserved G-hairpin. The black triangle indicates the conserved zinc-binding motif. GAS2 (mouse):

P11862, GAS2 (human): O43903, VAB-10 (elegans): G5EFM3, Shot (drosophila): Q7KJN8, MACF1 (mouse): Q9QXZ0, MACF1a (zebrafish): A0A8M3AL24, MACF1 (human): Q9UPN3, DST (human): Q03001, DST (mouse): Q91ZU6. **b** Overall analysis of the electrostatic potential of the contact surfaces showed complementary charges between the GAS2-GAR domain and the MT. The black circle shows the GAS2-GAR domain well bound to charge complementary concave of the intra-tubulin dimer. The cyan box shows negatively charged amino acid D163 in β-tubulin. The lime box shows the positive charged amino acid D163 in α-tubulin. **c** Bottom of the GAR domain is full of positively charged amino acids.
