## Supplementary material for "Dimerization GAS2 mediates microtubule and F-actin crosslinking": Description of Additional Video Files

**Video1 2μM Bare MT**

**Description:** Dark-field were used to observe bare MT. Taxol-stabilized MTs (2 μM) samples were imaged using the dark-field microscopy (BX53, Olympus). Bare MT is disordered as individual filaments. The scale bar represents 20 μm.

**Video2 1μM GAS2-FL bundle MT**

**Description:** Dark-field were used to observe MT bundling induced by GAS2-FL. Taxol-stabilized MTs (2 μM) were incubated with 1.0 μM GAS2-FL proteins in BRB80 at room temperature for 30 min. The samples were then imaged using dark-field microscopy (BX53, Olympus). MTs formed brightness bundling array cluster. The scale bar represents 20 μm.

**Video3 1μM GAS2ΔC couldn't bundle MT**

**Description:** Dark-field were used to observe MT bundling induced by GAS2ΔC. Taxol-stabilized MTs (2 μM) were incubated with 1.0 μM GAS2ΔC proteins in BRB80 at room temperature for 30 min. The samples were then imaged using dark-field microscopy (BX53, Olympus). MTs were still disordered as individual filaments. The scale bar represents 20 μm.

**Video4 1μM GAS2-GAR bundle MT**

**Description:** Dark-field were used to observe MT bundling induced by GAS2-GAR. Taxol-stabilized MTs (2 μM) were incubated with 1.0 μM GAS2-GAR proteins in BRB80 at room temperature for 30 min. The samples were then imaged using dark-field microscopy (BX53, Olympus). MTs formed brightness bundling array cluster. The scale bar represents 20 μm.

**Video5 1 μM GAS2-GARΔC couldn't bundle MT**

**Description:** Dark-field were used to observe MT bundling induced by GAS2-GARΔC. Taxol-stabilized MTs (2 μM) were incubated with 1.0 μM GAS2-GARΔC proteins in BRB80 at room temperature for 30 min. The samples were then imaged using dark-field microscopy (BX53, Olympus). MTs were still disordered as individual filaments. The scale bar represents 20 μm.
